# Control of N_2_ fixation and NH_3_ excretion in *Azorhizobium caulinodans* ORS571

**DOI:** 10.1101/2022.04.13.488174

**Authors:** Timothy L Haskett, Ramakrishnan Karunakaran, Marcelo Bueno Batista, Ray Dixon, Philip S Poole

**Affiliations:** Department of Plant Sciences, University of Oxford, Oxford, OX1 3RB, UK; Department of Molecular Microbiology, John Innes Centre, Norwich, NR4 7UH, UK

## Abstract

Due to the costly energy demands of N_2_ fixation, diazotrophic bacteria have evolved complex regulatory networks that permit expression of the N_2_-fixing catalyst nitrogenase only under conditions of N starvation, whereas the same condition stimulates upregulation of high-affinity NH_3_ assimilation by glutamine synthetase (GS), preventing excess release of excess NH_3_ for plants. Diazotrophic bacteria can be engineered to excrete NH_3_ by interference with GS, however control is required to minimise growth penalties and prevent unintended provision of NH_3_ to non-target plants. Here, we attempted two strategies to control GS regulation and NH_3_ excretion in our model cereal symbiont *Azorhizobium caulinodans Ac*LP, a derivative of ORS571. We first attempted to recapitulate previous work where mutation of both P_II_ homologues *glnB* and *glnK* stimulated GS shutdown but found that one of these genes was essential for growth. Secondly, we expressed unidirectional adenylyltransferases (uATs) in a Δ*glnE* mutant of *Ac*LP which permitted strong GS shutdown and excretion of NH_3_ derived from N_2_ fixation and completely alleviated negative feedback regulation on nitrogenase expression. We placed a *uAT* allele under control of the NifA-dependent promoter P*nifH*, permitting GS shutdown and NH_3_ excretion specifically under microaerobic conditions, the same cue that initiates N_2_ fixation, then deleted *nifA* and transferred a rhizopine-inducible *nifA*_*L94Q/D95Q*_*-rpoN* controller plasmid into this strain, permitting coupled rhizopine-dependent activation of N_2_ fixation with NH_3_ excretion. In future, this highly sophisticated and multi-layered control circuitry could be used to activate N_2_ fixation and NH_3_ excretion specifically by *Ac*LP colonising transgenic rhizopine producing cereals, targeting delivery of fixed N to the crop, and preventing interaction with non-target plants.

**Author Summary:** Inoculation of cereal crops with associative “diazotrophic” bacteria that convert atmospheric N_2_ to NH_3_ could be used to sustainably improve delivery of nitrogen in agriculture. However, due to the costly energy demands of N_2_ fixation, natural diazotrophic bacteria have evolved to conserve energy by preventing excess production of NH_3_ and release to the plants. Diazotrophs can be engineered for excess NH_3_ production and release, however genetic control is required to minimise growth penalties and prevent unintended provision of NH_3_ to non-target weed species. Here, we engineer control of N_2_ fixation and NH_3_ excretion in response to the signalling molecule rhizopine which is produced by transgenic barley. This control could be used to establish plant host-specific activation of N_2_ fixation and NH_3_ release following root colonisation in the field, minimising bacterial energy requirements in the bulk soil and preventing provision of NH_3_ to non-target plants.

## Introduction

Nitrogen (N) is an essential constituent of all biological organisms, but metabolically accessible forms are scarce in most environments, restricting biomass production. In agriculture, productivity of cereal crops, which are a staple of human dietary requirements, requires large-scale supplementation with synthetic N fertilisers to meet global food security requirements (1). However, synthesis and excessive application of N fertilisers has a large energy cost, causes CO_2_ release and results in loss of reduced N to the environment, which has doubled reactive N in the atmosphere and polluted waterways causing eutrophication and O_2_-depleted dead zones (2). In contrast, N fertilisers are largely unaffordable to small-hold farmers in developing countries such as those in Sub-Saharan Africa (3), restricting yields to a fraction of their maximum potential (4). Inoculation of cereals with root-associative diazotrophic bacteria that convert atmospheric N_2_ gas to NH_3_ through the action of O_2_-labile nitrogenase represents an affordable and sustainable alternative to the use of N fertilisers in agriculture (5–7). Although associative diazotrophs have been estimated to fix up to 70 kg N ha^−1^ year^−1^ in agricultural systems (8), responses to inoculation are typically inconsistent due to sub-optimal competitiveness for root colonisation and persistence in soil (9–12). Furthermore, due to the costly energy demands of N_2_ fixation, which consumes at least 16□mol ATP per mol N_2_ fixed *in vitro*, bacteria have evolved complex regulatory networks that permit expression and activity of the N_2_-fixing catalyst nitrogenase only under conditions of N starvation, whereas the same condition stimulates upregulation of high-affinity NH_3_ assimilation by glutamine synthetase (*glnA*, GS), preventing excess release of excess NH_3_ for plants (13, 14).

Associative diazotrophic bacteria can been engineered for excess production and excretion of NH_3_ by several strategies (13, 15, 16). For example, in *Azotobacter vinelandii*, insertional inactivation of *nifL*, which encodes an O_2_ as well as N and C sensing anti-activator of the nitrogenase master regulator NifA, drives constitutive nitrogenase activity resulting in excretion of NH_3_ from the cell (17–20). The same effect was achieved by expressing mutant *nifA* alleles that are resistant to inhibition by NifL (18, 21, 22). While excess NH_3_ production itself is likely to activate regulatory feedback mechanisms reducing GS biosynthetic activity and NH_3_ assimilation (15), mutating *glnA* (23–27) or genes involved in GS regulation may also be required to inhibit NH_3_ assimilation more strongly and favour NH_3_ excretion (28, 29).

Bacterial GS belongs to the “class I” type enzymes comprised of 12 identical subunits which are each adenylylated or de-adenylated by a bidirectional adenylyl transferase (AT, encoded by *glnE*) at the Tyr_397_ residue, with the fully de-adenylylated GS form being biosynthetically active and vice versa (30). Directionality of the ATase reaction is regulated by the post-translational modification state of P_II_ signal transduction proteins (31). The activity of P_II_ proteins is regulated by uridylylation/deuridylylation by the bidirectional uridylyltransferase (UT) GlnD which represents the most basal regulator in the cascade and can directly sense N status of the cell (32). GlnD uridylylates P_II_ under conditions of N-starvation and the resulting P_II_-UMP ultimately triggers dephosphorylation of ATase and hence deadenylylation and activation of GS (33). In *Azorhizobium caulinodans* (*Ac*), insertional inactivation of both P_II_ homologues *glnB* and *glnK* produced a mutant that was unable to activate GS by deadenylylation, driving NH_3_-insensitive N_2_ fixation and excretion of NH_3_ into the growth media (28). Critically, this engineering strategy does not appear to be universally applicable as P_II_ is essential for NifA and nitrogenase activity in some bacteria (34, 35), whereas it is essential for growth in others (36, 37). In a Δ*glnE* ATase mutant of *Azospirillum brasilense*, complementation with unidirectional adeyltransferase (uAT) alleles that encoded only the C-terminal adenylylation domain (31) drove strong adenylylation of GS resulting in excretion of NH_3_ into the growth media (29). This strategy likely represents a more universally applicable approach for engineering NH_3_ excretion in diazotrophs because the ATase is highly conserved, has a specific function, and can be readily mutated across diverse diazotrophic bacterial taxa (15, 38–40), albeit the mutation appears to be lethal in the heterotroph *Mycobacterium tuberculosis* (41, 42).

From an agricultural perspective, there are three major caveats of engineering diazotrophic bacteria for excessive production and excretion of NH_3_; i) uncontrolled *nifA* and(or) nitrogenase expression has a severe energy burden on the cell that could abolish competitiveness for root colonisation; ii) interference with GS activity typically renders strains auxotrophic for the essential amino acid glutamine, which could further reduce competitiveness; and iii) NH_3_ excreting bacteria have potential to supply NH_3_ to non-target weed species following promiscuous colonisation in the field. Therefore, establishing control of N_2_ fixation and NH_3_ excretion will be crucial for the optimisation of strains as agricultural inoculants. Control of NH_3_ excretion has already been achieved in *A. vinelandii* by establishing IPTG-dependent expression of *glnA* (27), and in *A. brasilense* by establishing anhydro-tetracycline inducible expression of uATs (29, 43). However, use of plant-derived signals to control N_2_-fixation and NH_3_ excretion would be far more applicable in the environment and could impart partner-specificity to target delivery of fixed N to crops and prevent interactions with non-target host plants following promiscuous colonisation (44, 45).

We previously developed synthetic rhizopine signalling between barley and the model endophyte *Azorhizobium caulinodans Ac*LP that stimulates transcriptional activation of the mutant nitrogenase master regulator *nifA*_*L94Q/D95Q*,_ which partially escapes nitrogen regulation, and when paired with the sigma factor RpoN drives N_2_ fixation in bacteria colonising rhizopine producing (*RhiP*) barley roots (44, 46, 47). Here, we demonstrate that wild-type and engineered *Ac* strains do not release fixed N as NH_3_ into the growth media when cultured under N_2_-fixing conditions and therefore sought to engineer this trait by interfering with high-affinity NH_3_ assimilation catalysed by GS. In our attempts to recapitulate NH_3_ excreting *glnB glnK* double mutants of *Ac*LP (28), we found that deletion of both P_II_ homologues was only possible when second copy of *glnB* was first integrated into the chromosome suggesting one of the P_II_ homologues were essential for growth. GS and nitrogenase activity in the resulting strain exhibited minimal variation from that of the wild-type, but nevertheless the strain excreted low levels of NH_3_ into the growth media. To optimise rates of NH_3_ excretion, we utilised a second engineering strategy where a *Ac*LPΔ*glnE* mutant was complemented with uATs. In congruency with similar experiments performed in *A. brasilense* (29), uAT expression drove strong shutdown of GS, but also completely alleviated negative feedback inhibition of nitrogenase by NH_3_ and stimulated NH_3_ excretion. By placing uAT expression under control of NifA, we established control of these traits by the same cues that initiate N_2_ fixation, then transferred rhizopine control of *nifA*_*L94Q/D95Q-*_*rpoN* into this strain linking activation of N_2_-fixation and NH_3_ excretion (Fig 1). This highly sophisticated control circuitry represents a significant milestone in the development of a “synthetic symbiosis” where N_2_ fixation and NH_3_ excretion could be activated in bacteria specifically colonising target cereals, targeting delivery of N to the crops while avoiding potential interactions with non-target plants.

**Figure 1.**
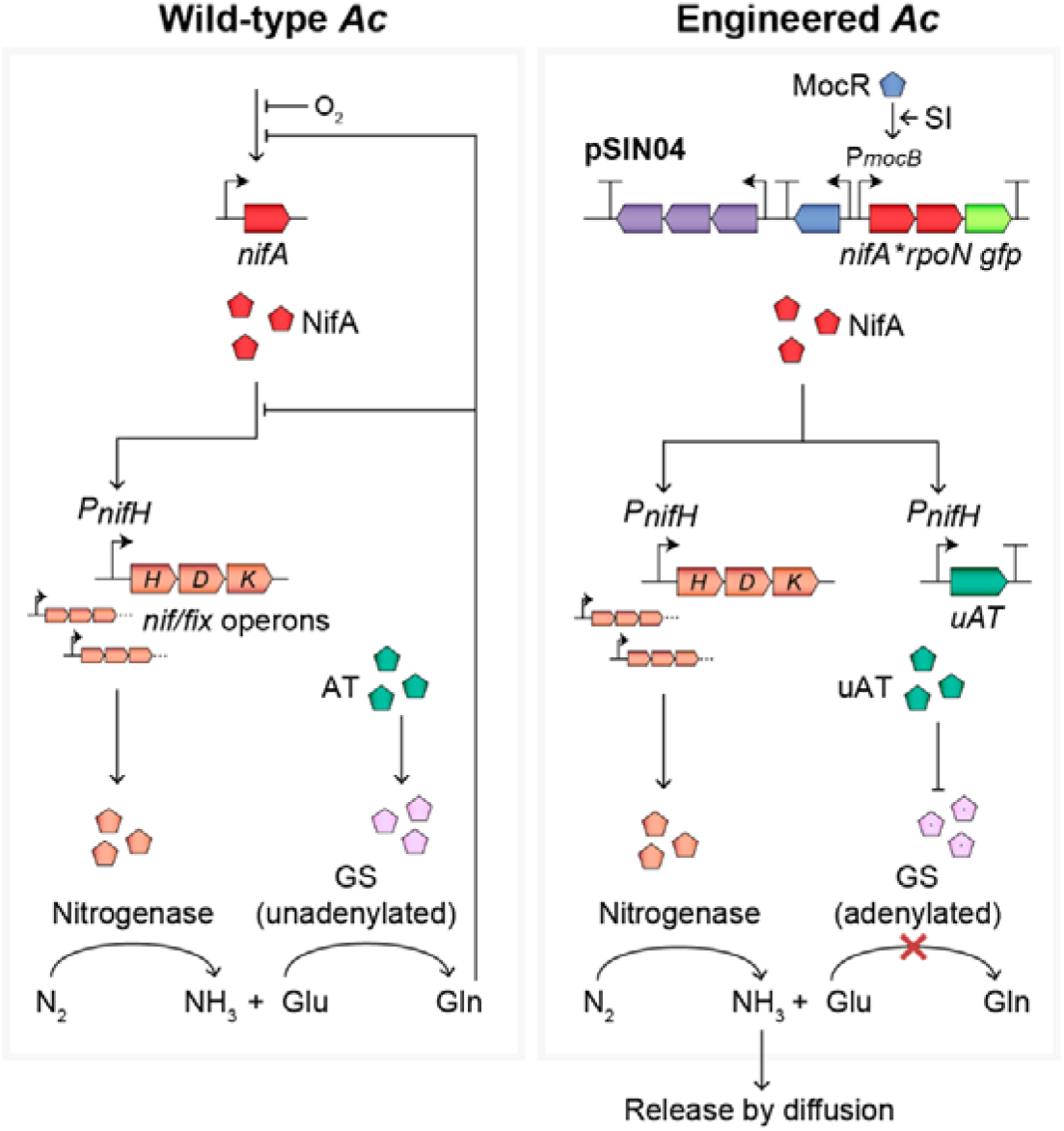
Model for rhizopine control of N_2_ fixation and NH_3_ excretion in *Ac*. In the wild-type bacterium, NifA is activated under N_2_-fixing (N-free microaerobic) conditions leading to transcription of nitrogenase (*nif* and *fix*) genes and subsequently N_2_ fixation. Under the same conditions, P_II_-UMP stimulates the adenylyltransferase (AT) to activate glutamine synthetase (GS) by deadenylylation. GS catalyses assimilation of NH_3_ via the conversion of glutamate (Glu) to glutamine (Gln), which feeds back to repress *nif* expression and NifA activity, preventing excess production and release of NH_3_ from the cell. In our engineered strain *Ac*PU-R22 carrying pSIN04, *nifA*_*L94Q/F95Q*_ is expressed by addition of rhizopine to the culture, driving nitrogenase expression and N_2_ fixation. Additionally, NifA induces transcription of the unidirectional adenylyltransferase (uAT) which drives shutdown of GS by adenylylation, preventing NH_3_ assimilation. Because shutdown of GS prevents glutamine biosynthesis, repression on *nif* gene expression and NifA activity is alleviated, resulting in excess production of NH_3_ and release from the cell.

## Results

### Deletion or strong repression of the *P*_*II*_ genes is lethal

It was previously demonstrated that insertional inactivation of the *Ac P*_*II*_ genes *glnB* and *glnK* stimulates shutdown of GS by adenylylation and alleviates negative feedback inhibition of nitrogenase by the product NH_3_, preventing NH_3_ assimilation and favouring excretion into the growth media (28). We attempted to recapitulate these experiments in *Ac*LP, a derivative of *Ac* harbouring a mini-Tn7 *attB* integration site stably recombined into its chromosome, by constructing a markerless deletion of *glnB* and replacing *glnK* with an omega (Ω)-spectinomycin resistance (Sp) cassette.

Although the single Δ*glnB* and Δ*glnK*::ΩSp mutations were readily acquired, we were unable to acquire the double mutant by introduction of the Δ*glnK*::ΩSp mutation into *Ac*LPΔ*glnB* when selection was performed on rich or minimal media supplemented with glutamine as a sole N source, suggesting the resulting phenotype was lethal. To explore this notion further, we integrated into the chromosome of *Ac*LPΔ*glnB* a construct encoding *glnB* with the strong ribosome binding site (RBS) RStd expressed from the IPTG de-repressible promoter P*lac* (Fig 2a) and were subsequently able to acquire the Δ*glnB* Δ*glnK*::ΩSp double mutation when selection was performed on rich media in the absence of IPTG, confirming that one of the P_II_ proteins was essential for growth.

**Figure 2.**
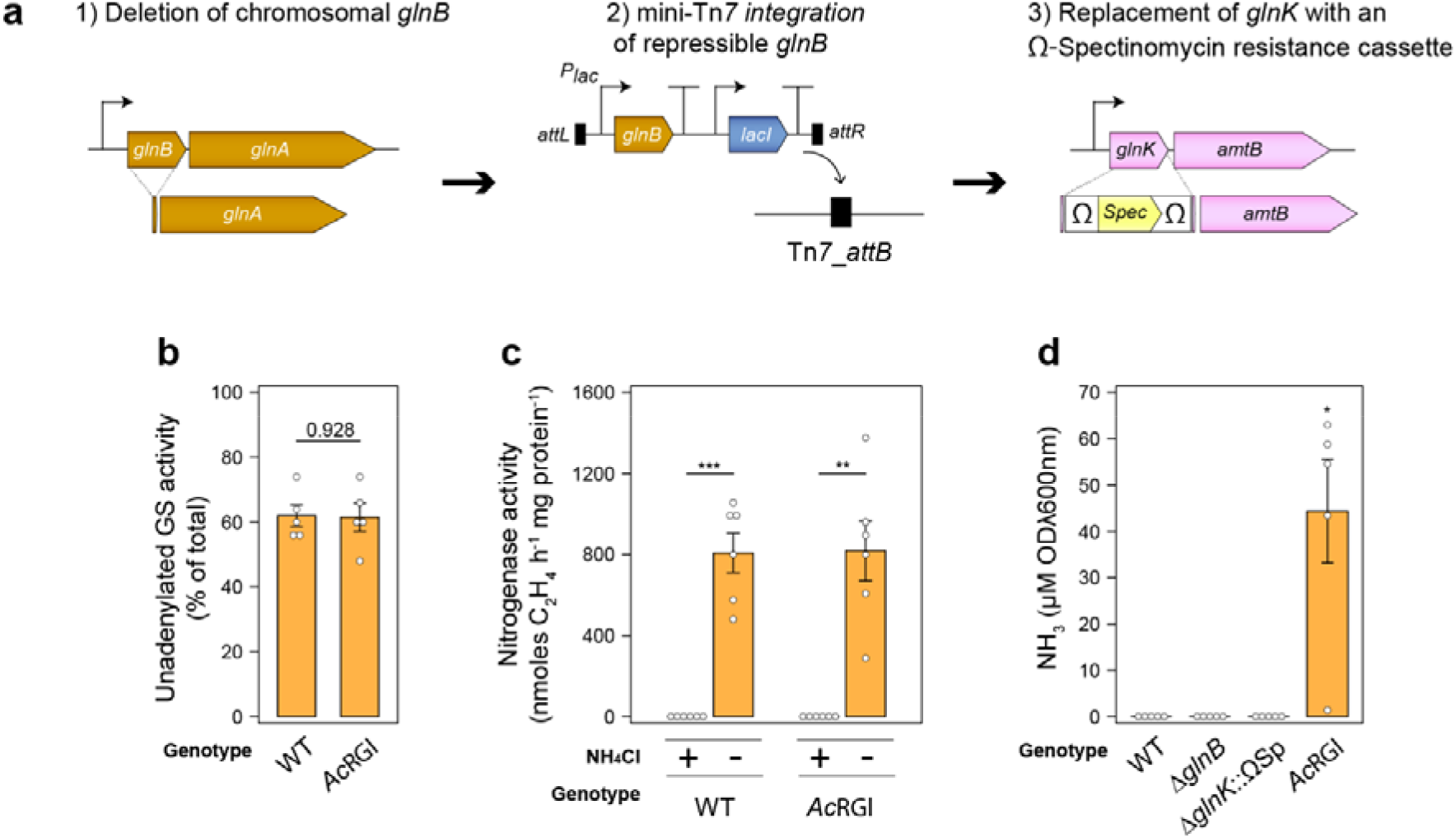
Strong repression of *glnB* in a *glnK* mutant has minimal effect on glutamine synthetase and nitrogenase activity but drives low-level NH_3_ excretion. **(a)** Strategy for generating strain *Ac*RGl with the double Δ*glnB* and Δ*glnK*::ΩSp mutation following integration of an IPTG-derepressible *glnB* gene into the chromosome of *Ac*LP. **(b)** Activity of the unadenylylated (active) form of GS in *n = 5* wild-type (WT) or *Ac*RGl cultures incubated for 24-h as determined by γ-glutamyl transferase assays in the presence or absence of 60 mM MgCl_2_ (see S2 Fig for total activity). **(c)** Nitrogenase activity measured by acetylene reduction in *n = 6* cultures between 3-h – 21-h **(d)** Spectrophotometric determination of NH_3_ in media of *n = 5* cultures grown for 24-h. Cultures for all assays were grown in N_2_-fixing conditions (N-free UMS media with 3% O_2_ in the headspace). Error bars represent one SEM. Independent two-tailed students t-tests were used to compare means. Exact P-values are provided where P > 0.05. *P < 0.05, **P < 0.01, ***P < 0.001. The wild-type *Ac*LP was used as a reference group for comparison of means in panel (e).

We next sought to test whether reduced translation of the introduced *glnB* module would stimulate GS shutdown and NH_3_ excretion by tuning the ribosome binding site (RBS). Seven synthetic RBS’ were experimentally demonstrated to produce translation rates spanning two to three orders of magnitude (S1 Fig), but only when *glnB* fused to the strongest RBS “RStd” and integrated into the *Ac*LPΔ*glnB* chromosome were we able to subsequently isolate the Δ*glnK*::ΩSp mutation (hereby termed strain *Ac*RGl), suggesting that *glnB* had been repressed as much as was tolerable. We assessed total GS specific activity and that of the unadenylylated active enzyme in *Ac*RGl by performing γ-glutamyl transferase assays on whole cells in the presence or absence of 60 mM MgCl_2_ which specifically inhibits the adenylylated enzyme (48), and found that mutant exhibited higher total GS activity compared to the wild-type (S2a Fig), presumably due to elevated *glnA* expression (S2b Fig) as is typical of *glnB* mutants (25, 28), whereas the adenylylation state of GS (depicted here as percentage of unadenylylated GS activity) was unchanged (Fig 2b). We also found that the specific nitrogenase activity of strain *Ac*RGl was no different from that of the wild-type, being repressed by supplementation of 10 mM NH_3_Cl into the growth media (Fig 2c). Spectrophotometric quantification of NH_3_ was next performed using the indophenol method (49) on the strains grown for 24-h under N_2_-fixing conditions (here defined as N-free UMS with O_2_ in the headspace adjusted to 3%). No NH_3_ was detected in the wild-type or Δ*glnB* and Δ*glnK*::ΩSp single mutants, whereas we detected trace amounts of NH_3_ in the growth media of strain *Ac*RGl (Fig 2d). Given that construction of the *Ac*LPΔ*glnB* Δ*glnK*::ΩSp double mutant was lethal, and strong repression of *glnB* had minimal effect on GS and nitrogenase regulation permitting only low-level NH_3_ excretion, we concluded that these strategies were inadequate to establish control of NH_3_ excretion in *Ac*LP and opted to pursue an alternative strategy.

### uAT expression drives GS inactivation and NH_3_ excretion

In a Δ*glnE* mutant of *A. brasilense*, controlled expression of a N-terminal truncated AT consisting of only the adenylylation domain results in unidirectional activity driving strong inactivation of GS by adenylylation and excretion of NH_3_ into the growth media (29). We recapitulated these experiments in a Δ*glnE* mutant of *Ac*LP by using the *Sinorhizobium meliloti* derived P*nodA* promoter (S3 Fig), to drive expression of a series of truncated uATs derived from *Ac* or those previously described for *E. coli* (Fig 3a-b) (29). We assessed GS specific activity and that of the unadenylylated enzyme using γ-glutamyl transferase assays on cells grown in N_2_-fixing conditions for 3-h and confirmed that leaky non-induced uAT expression stimulated GS adenylylation (Fig 3c), while having minimal effect on total GS specific activity relative to wild-type bacteria (S4 Fig). The strains also excreted between 0.1-1.5mM of NH_3_ after 24-h incubation in N_2_-fixing conditions, whereas the wild-type and Δ*glnE* mutant did not excrete detectable levels of NH_3_ (Fig 3d). Interestingly, we found that NH_3_ excretion was sub-optimal when the P*nodA* promoter controlling *uAT* expression was induced with saturating levels (5 uM) of naringenin (S5 Fig), suggesting that *uAT* overexpression is metabolically detrimental, as was observed in *A. brasilense* (43). This indicated that more finely tuned *uAT* expression would be critical to achieve stringent control of GS adenylylation in *Ac*LP.

**Figure 3.**
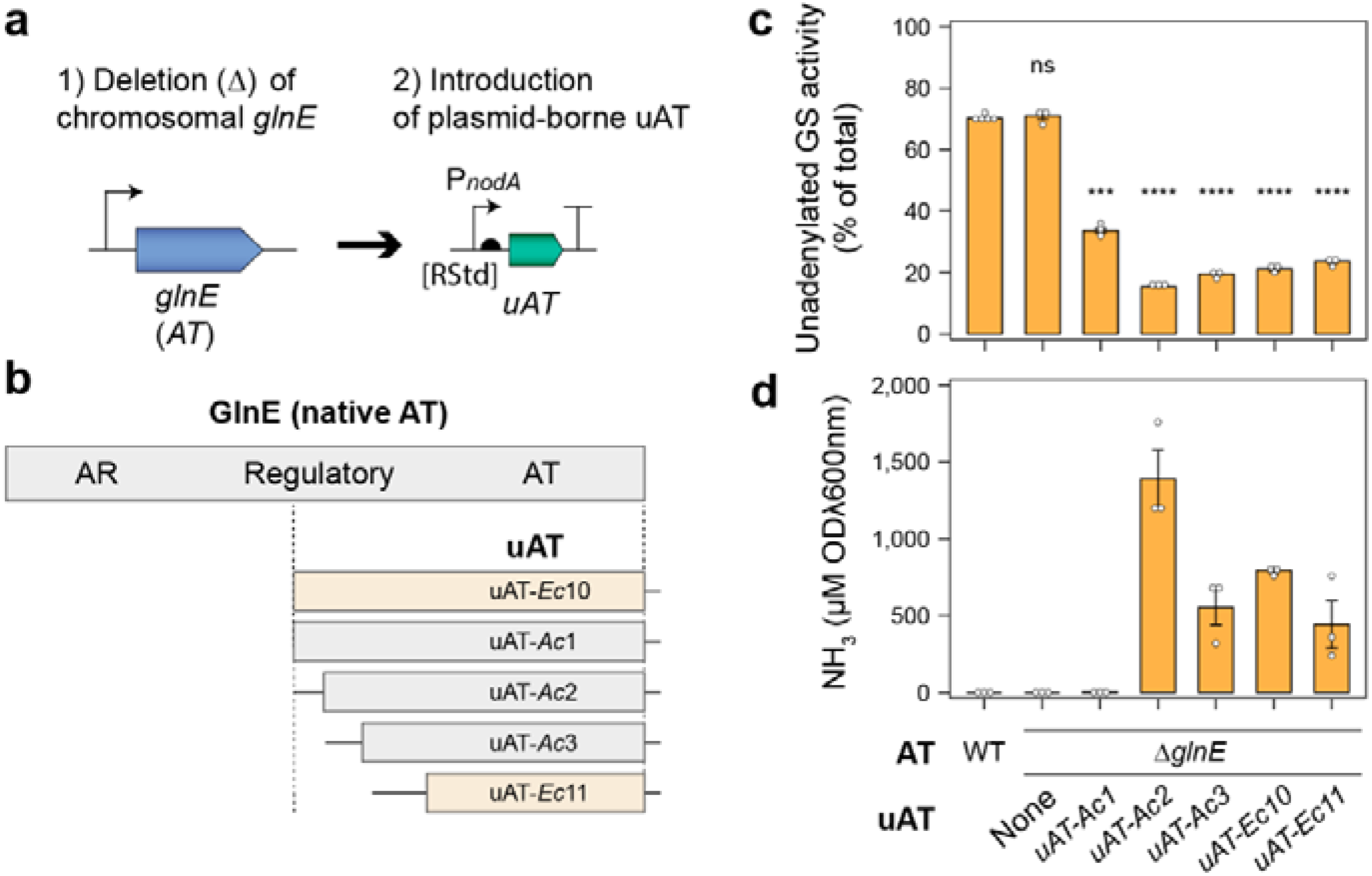
uAT expression drives GS adenylylation and NH_3_ excretion in a Δ*glnE* background. **(a)** Strategy for complementation of the Δ*glnE* mutation with naringenin-inducible unidirectional adenylyltransferases (uAT) expressed from low-copy parent plasmid pOPS1536. **(b)** A series of truncated uAT proteins were used in this study. The *uAT-Ec10* and *uAT-Ec11* alleles are derived from *E. coli* were described previously (29), whereas *uAT-Ac* alleles are derived from *Ac*LP. The nucleotide sequences for these alleles are provided in S1 File. **(c)** Activity of the unadenylylated (active) form of GS in *n = 5* for *Ac*LP (wild-type, WT) or *n* = 3 cultures incubated for 3-h in N_2_-fixing conditions (N-free UMS media with 3% O_2_ in the headspace) without the inducer naringenin as determined by γ-glutamyl transferase assays in the presence or absence of 60 mM MgCl_2_ (see S4 Fig for total activity). **(d)** Spectrophotometric determination of NH_3_ in media of *n =* 3 cultures grown for 24-h in N_2_-fixing conditions. Error bars represent one SEM. Independent two-tailed students t-tests with the Bonferroni-holm adjustment were used to compare means using the wild-type *Ac*LP as a reference group. Not significant (ns P > 0.05), ***P < 0.001, ****P < 0.0001.

### Shutdown of glutamine biosynthesis alleviates negative feedback on nitrogenase

Expression of *uAT* restricts glutamine production via the high affinity GS-dependent NH_3_ assimilation pathway, providing us with a unique opportunity to tease apart the effects of NH_3_ and glutamine on the nitrogenase (*nif*) gene expression. We postulated that NH_3_ must first be converted into glutamine to mediate repression of *nif* genes and tested this hypothesis first by examining expression of P*nifH* fused to *GFP* on plasmid pOPS1213 in wild-type bacteria and in *Ac*LPΔ*glnE* expressing the *uAT*-*Ac2* allele from the non-induced P*nodA* promoter on a second plasmid. As expected, P*nifH*::*GFP* activity in both strains grown under microaerobic conditions (3% O_2_ in the headspace) was strongly repressed by supplementation with 10 mM glutamine however, while P*nifH*::*GFP* was repressed in the wild-type by supplementation with 10 mM NH_4_Cl, P*nifH*::*GFP* expression was not repressed by NH_4_Cl in *Ac*LPΔ*glnE* expressing *uAT-Ac2* (Fig 4a). We observed a similar pattern when nitrogenase activity was assessed by ARAs (Fig 4b-c), indicating that NH_3_ itself has no effect on negative feedback regulation of *nif* genes but must be converted into glutamine or potentially other amino acids to facilitate repression. Engineering NH_3_ excreting bacteria by targeted GS shutdown therefore has two advantages; i) alleviating negative feedback regulation of *nif* genes and ii) preventing NH_3_ assimilation to favour release.

**Figure 4.**
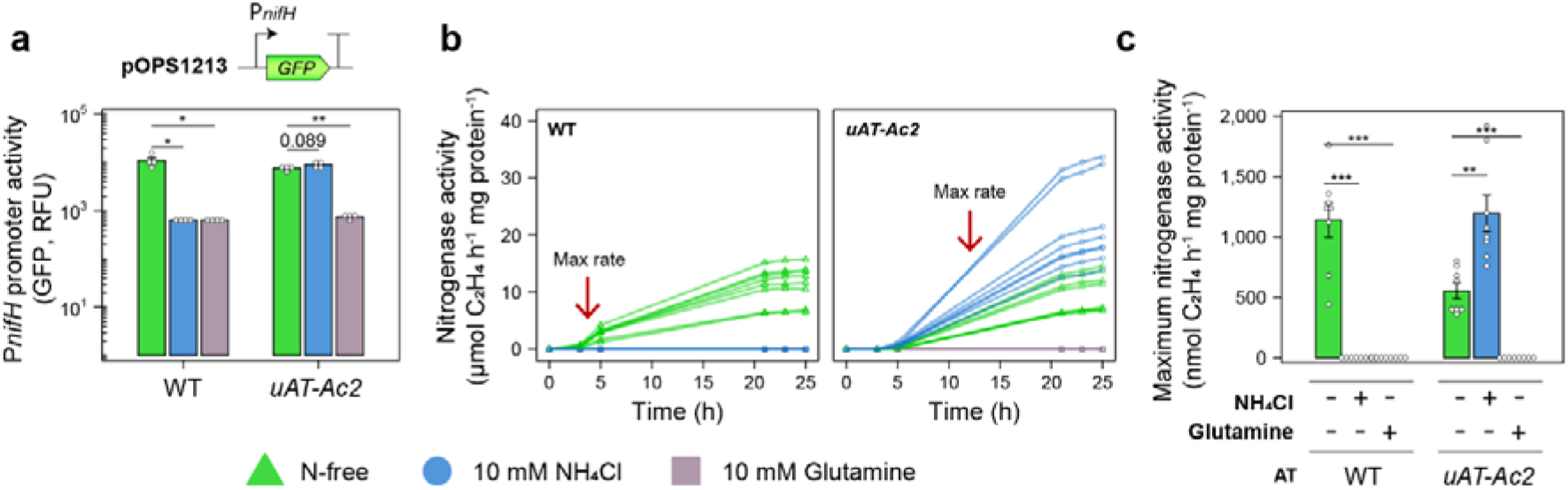
uAT expression abolishes negative feedback regulation on nitrogenase. **(a)** A P*nifH*::*GFP* reporter carried on plasmid pOPS1213 was mobilised into the wild-type (WT) *Ac*LP and *Ac*LPΔ*glnE* expressing *uAT-Ac2* on a second low-copy plasmid and induction was measured in *n =* 4 cultures grown for 24-h under the conditions indicated. Relative fluorescence units (RFU) are defined here as GFP fluorescence / OD600λnm **(b)** Nitrogenase activity was measured by acetylene reduction in *n =* 8 cultures grown under N_2_-fixing conditions (N-free UMS media with 3% O_2_ in the headspace) and **(c)** the maximum rates are presented. Error bars represent one SEM. Independent two-tailed students t-tests with Bonferroni-holm adjustment were used to compare means. Exact P-values are provided where P > 0.05. **P < 0.01, ***P < 0.001. WT bacteria grown at 3% O_2_ in N-free conditions was used as a reference group for statistical comparisons in panel (a).

### NifA control of *uAT* expression

As a direct consequence of engineering NH_3_ excretion through GS interference, bacteria typically become auxotrophic for glutamine. While this may not be non-problematic for cultures grown *in vitro* under gnotobiotic conditions, glutamine auxotrophs in the field would be unable to compete or persist in the soil and rhizosphere. In rhizobia-legume symbioses, rhizobia only restrict NH_3_ assimilation after infecting the low-oxygen environment of the nodule and differentiating into an N_2_ fixing bacteroid (50, 51), allowing them to maintain competitiveness during their free-living state in the soil. To mimic this regulation, we fused the *uAT-Ac2* allele to native or synthetic RBSs and placed these under control of the NifA-inducible P*nifH* promoter on mini-Tn*7* delivery plasmids, then integrated these into the chromosome of *Ac*LPΔ*glnE*, creating strains *Ac*PU-RStd, *Ac*PU-R1, *Ac*PU-R22, *Ac*PU-R31, *Ac*PU-Rnat and *Ac*PU-R28 (Fig 5a). When grown under aerobic (21% O_2_) conditions in the presence of 10 mM NH_4_Cl, growth of *Ac*LPΔ*glnE* expressing the *uAT*-*Ac2* allele from the non-induced P*nodA* promoter was almost entirely abolished compared to where glutamine was provided as a source of N (Fig 5b and S6 Fig). In contrast, the growth characteristics of strains expressing *uAT-Ac2* from the P*nifH* promoter were reminiscent of the wild-type *Ac*, except for strains where *uAT-Ac2* was fused to the strongest RBS’ RStd or R1, which increased mean generation times (MGT) but did not affect the total biomass at stationary phase (Fig 5b and S6 Fig). We next assessed GS regulation by γ-glutamyltransferase assays and confirmed that under aerobic conditions in the presence of 10 mM NH_4_Cl, the percentage of active deadenylylated GS activity in strains *Ac*PU-R1, *Ac*PU-R22, and *Ac*PU-R3 closely resembled that of the wild-type, suggesting that NH_3_ assimilation was functional. When grown under microaerobic conditions (3% O_2_) in the presence or absence of 10 mM NH_4_Cl, GS in wild-type *Ac*LP was activated by de-adenylylation, whereas GS in all *Ac*LPΔ*glnE* strains expressing uAT-*Ac*2 from the P*nifH* promoter became more heavily inactivated by adenylylation under the same conditions (Fig 5c), with the percentage unadenylated GS activity correlating negatively with the strength of RBS fused to *uAT-Ac2*. We finally performed NH_3_ excretion assays on the engineered strains and found that each excreted NH_3_ into the growth media after 24-h, except for where *uAT-Ac2* was fused to the weakest RBS [R28] (Fig 5d). Overall, the data suggested that by expressing *uATs* from the P*nifH* promoter, GS shutdown can be controlled in response to the same cues that initiates N_2_ fixation.

**Figure 5.**
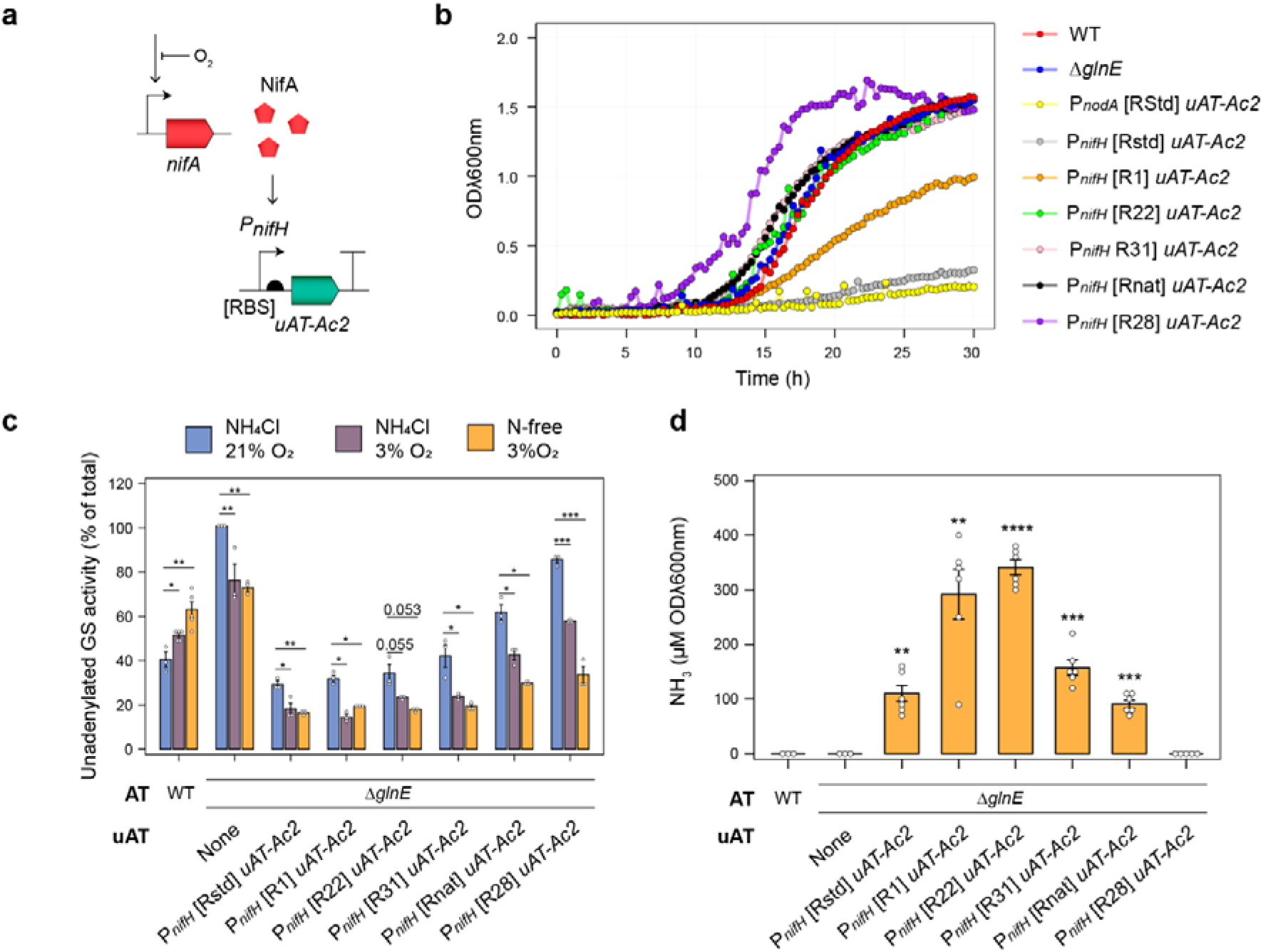
Coupled activation of N_2_ fixation and GS adenylylation via NifA-dependent expression of *uAT*. **(a)** Strategy for complementation of the Δ*glnE* mutation with NifA-inducible unidirectional adenylyltransferases (uAT) integrated into the chromosome using mini-Tn*7*. **(b)** Growth of control strains and those expressing uATs in UMS media supplemented with 20 mM succinate and either 10 mM glutamine or 10 mM NH_3_Cl under aerobic conditions. See S6 Fig for full growth statistics. **(c)** Activity of the unadenylylated (active) form of GS in *n =* 5 for wild-type (WT) *Ac*LP or *n* = 3 cultures incubated for 24-h in as determined by γ-glutamyl transferase assays in the presence or absence of 60 mM MgCl_2_. (**d**) Spectrophotometric determination of NH_3_ in media of *n =* 3 WT and Δ*glnE* or *n* = 5 cultures grown for 24-h in N_2_-fixing conditions (N-free UMS media with 3% O_2_ in the headspace). Error bars represent one SEM. Independent two-tailed students t-tests with the Bonferroni-holm adjustment were used to compare means using the wild-type *Ac*LP as a reference group. Exact P-values are provided where P > 0.05. **P < 0.01, ***P < 0.001. WT bacteria were used as a reference group for statistical comparisons in panels (a) and (b).

### Rhizopine-dependent control of N_2_ fixation, GS adenylylation and NH_3_ excretion

While NifA-dependent expression of nitrogenase and *uAT-Ac2* in *Ac*Δ*glnE* drives N_2_ fixation and GS inactivation leading to NH_3_ excretion, the lack of plant host-specific signalling to drive these processes could permit bacteria to supply NH_3_ to target crops and non-target weed species alike. We previously used synthetic rhizopine signalling to establish control of a mutant *nifA* allele (encoding NifA_L94Q/D95Q_) and *rpoN* in *Ac*LPΔ*nifA* carrying plasmid pSIN02, which drove partially NH_3_-resistant activation of nitrogenase activity specifically by bacteria occupying the roots of transgenic *RhiP* barley (46). We performed NH_3_ excretion assays on *Ac*LPΔ*nifA* carrying pSIN02 and found that this strain did not secrete NH_3_ into the growth media (S7 Fig). Thus, we opted to establish rhizopine control of the *nifA*_*L94Q/D95Q*_*-rpoN* operon in our strain *Ac*PU-R22 where *uAT-Ac2* expression placed under control by NifA. We first tested in *Ac*LP, induction of a new rhizopine receiver plasmid pSIR03 which was derived from the high-copy rhizopine receiver pSIR03 but carried an RK2 replicon for low-copy maintenance. Using *GFP* induction assays, we demonstrated that pSIR03 (Fig 6a) has a dynamic range of 162-fold in response to the rhizopine *scyllo-*inosamine (SI) and was induced in 93.08 ± [SEM] 0.32% of cells in populations when 10 μM SI was supplemented *in vitro* (Fig 6b-c & Table S1). We deleted the native *nifA* gene from strain *Ac*PU-R22 and introduced a rhizopine *nifA*_*L94Q/D95Q*_*-rpoN* controller plasmid pSIN04 which was derived from pSIR03 (Fig 6d). Expression of *nifA*_*L94Q/D95Q*_*-rpoN* under microaerobic conditions by addition of 10 μM SI into the media resulted in tightly controlled activation of nitrogenase that was unimpeded by addition of 10 mM of NH_3_ (Fig 6e). Moreover, GS was strongly adenylylated by addition of 10 SI to the media in both aerobic and microaerobic conditions (Fig 6f). Because NifA in many diazotrophs is inactivated when cells are grown at 21% O_2_ (52), we subsequently tested O_2_ tolerance of our NifA_L94/D95Q_ mutant protein by inducing expression of *nifA*_*L94Q/D95Q*_*-rpoN* in *Ac*LPΔ*nifA* carrying pSIN03 with rhizopine and monitoring activation of the P*nifH::GFP* promoter fusion (S8a Fig). Interestingly, the NifA_L94/D95Q_ protein activated P*nifH::GFP* 13-fold ± [SEM] 1.5 and 98-fold ± [SEM] 2.8 under aerobic and microaerobic conditions, respectively (S8b Fig), suggesting that the protein is tolerant to oxygen.

**Figure 6.**
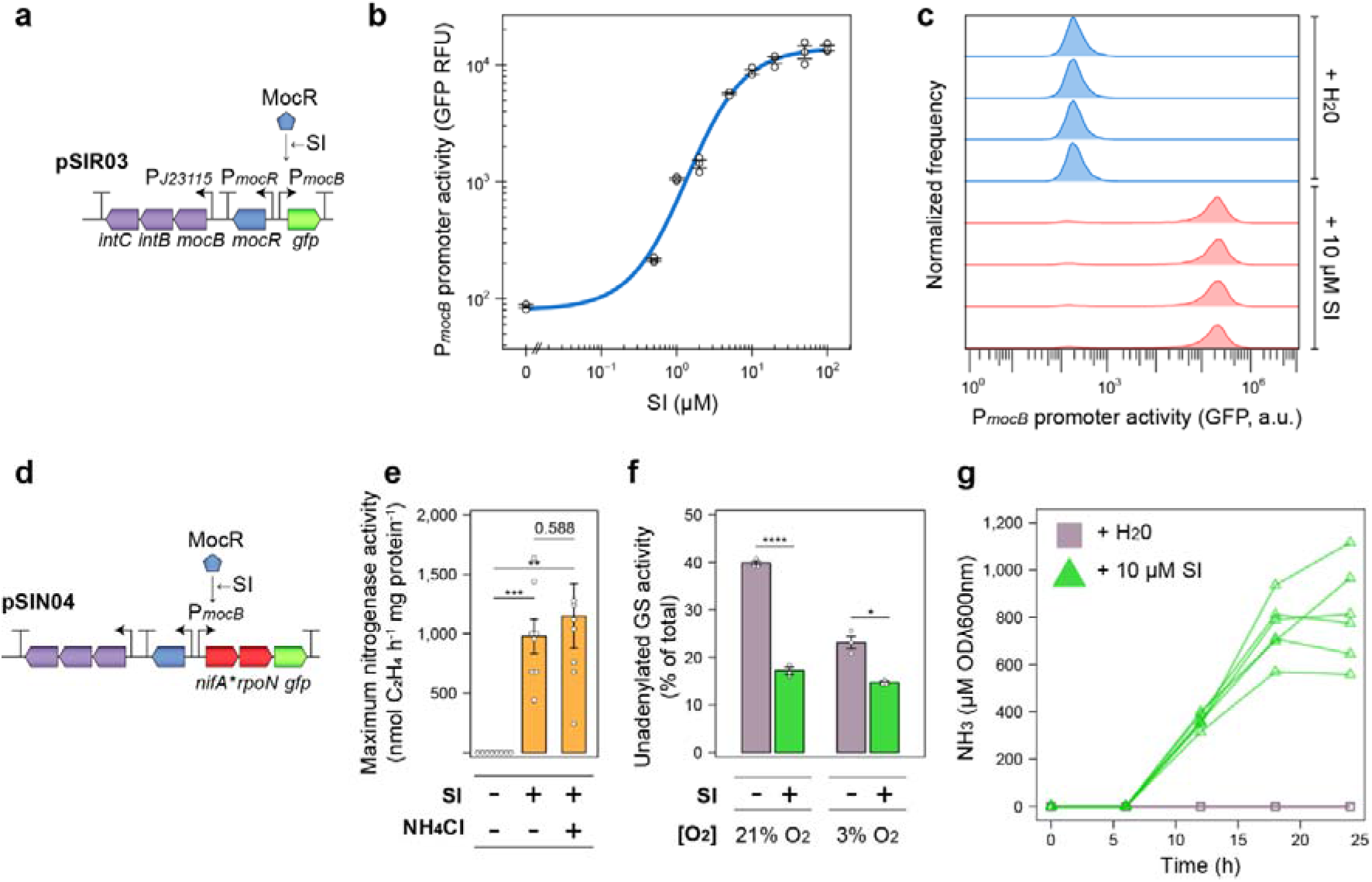
Rhizopine control of N_2_ fixation, GS adenylylation and NH_3_ excretion. **(a)** Genetic schematic (not to scale) of the low-copy (RK2 replicon) rhizopine receiver plasmid pSIR03. **(b)** Dose response of GFP induction in *Ac*LP (*n =* 3) harbouring pSIR03 with the rhizopine *scyllo*-inosamine (SI) supplemented *in vitro*. Relative fluorescence units (RFU) are defined here as GFP fluorescence / OD600λnm. **(c)** Flow-cytometry analysis of GFP fluorescence in *Ac*LP (*n =*4) harbouring pSIR03 incubated for 24-h in the absence or presence of 10 μM rhizopine. See Table S1 for full statistics. **(d)** Genetic schematic (not to scale) of the low-copy (RK2 replicon) rhizopine *nifA*_*L94Q/D95Q*_*-rpoN* controller plasmid pSIN04. **(e)** Optimal nitrogenase activity of *Ac*PU-R22 Δ*nifA* carrying pSIN04 measured between 5-h – 21-h by acetylene reduction in *n =* 6 cultures grown under microaerobic conditions (3% O_2_ in the headspace). **(f)** Activity of the unadenylylated (active) form of GS in *n* = 3 cultures of *Ac*PU-R22 Δ*nifA* carrying pSIN04 incubated for 24-h as determined by γ-glutamyl transferase assays in the presence or absence of 60 mM MgCl_2_. **(g)** Spectrophotometric determination of NH_3_ in media of *n =* 6 cultures of *Ac*PU-R22Δ*nifA* carrying pSIN04 grown in N_2_-fixing conditions (N-free UMS media with 3% O_2_ in the headspace) in the presence of absence of 10 μM SI. Error bars represent one SEM. Independent two-tailed students t-tests with the Bonferroni-holm adjustment were used to compare means. Exact P-values are provided where P > 0.5. **P < 0.01, ***P < 0.001, ****P < 0.0001.

We finally performed NH_3_ excretion assays on cultures of strain *Ac*PU-R22 carrying pSIN04 supplemented with 10 μM SI and demonstrated that the strain excreted 812.58 ± [SEM] 5.59 uM ODλ600nm^−1^ NH_3_ into the growth media after 24-h incubation in N_2_-fixing conditions at an optimal rate of 65.13 ± 7.35 uM OD600nm^−1^ h^−1^ (Fig 5g). Importantly, no NH_3_ excretion was detected in the absence of SI, demonstrating tight control. Overall, these experiments confirmed that we had established rhizopine control of N_2_-fixation, GS adenylylation and NH_3_ excretion in our engineered *Ac*LP strain.

## Discussion

In this study, we employed two strategies to interfere with GS and stimulate NH_3_ excretion in *Ac*LP. For our first strategy, we attempted to recapitulate previous experiments where insertional inactivation of the *P*_*II*_ genes *glnB* and *glnK* stimulated shutdown of GS by adenylylation and alleviated negative feedback inhibition of nitrogenase by the product NH_3_, preventing NH_3_ assimilation and favouring excretion into the growth media (28). Although we could delete either of the *glnB* or *glnK* genes from *Ac*LP, we were unable to delete both genes in the same strain unless a second copy of *glnB* was first introduced into the chromosome, suggesting at least one of the P_II_ coding sequences was essential for growth. Considering that a P*aph*::KIXX kanamycin resistance cassette was previously inserted to the 3’-end of the *Ac glnB* coding sequence (25) leaving most of the 5’-end reading frame intact, it seems possible that the GlnB protein may have retained some essential function unrelated to AT and GS activity. In contrast, previous insertion of the omega interposon into *Ac glnK* replaced a segment of the internal coding sequence and was therefore more likely to have abolished the function of the protein (53). Interestingly, similar *glnB* and *glnK* antibiotic cassette insertions have been made in the phototrophic diazotroph *Rhodobacter capsulatus*, resulting in NH_3_-insensitive NifA and nitrogenase expression and activity (54). However, attempts to delete both genes were also unsuccessful in this bacterium (55). Regardless of why deleting *glnB* and *glnK* is lethal, reproducing exact copies of the original *glnB* and *glnK* mutants (28) would likely be required to establish control of NH_3_ excretion in *Ac*LP, as we have shown here that deletion of *glnK* paired with strong repression of *glnB* had minimal effect on nitrogenase or GS regulation and permitted only low-level NH_3_ excretion.

As was previously demonstrated in *A. brasilense* (29), expression of *E. coli* or *Ac*-derived uATs in our *Ac*LPΔ*glnE* mutant resulted in strong GS shutdown and high rates of NH_3_ excretion when grown in N_2_-fixing conditions. We also found that while nitrogenase expression and activity is repressed in microaerobic NH_3_ or glutamine-fed cultures of *Ac*LP, shutdown of glutamine biosynthesis by uAT expression resulted in nitrogenase expression that was unimpeded by NH_3_ but still repressed by glutamine, suggesting that NH_3_ must first be converted to glutamine or potentially other amino acids such as asparagine (56) to facilitate repression. This same effect was previously reported in phototrophic *Anabaena* spp (57, 58) and *Rhodobacter sphaeroides* (59) where GS activity was shutdown using the chemical inhibitor L-Methionine sulfoximine, and in *Klebisella pneumoniae* mutants unable to grow on NH_3_ as a sole source of N (60). Moreover, In *R. capsulatus*, where N_2_-fixation is repressed in response to added NH_3_ at three levels; a) NtrC-dependent transcription of *nifA*; b) NifA-dependent transcription of nitrogenase; and c) DraT-DraG-dependent ADP ribosylation of nitrogenase (61, 62); all three levels of regulation were non-responsive to NH_3_ following shutdown of GS by insertional inactivation of both P_II_ genes (54). Thus, it seems plausible that shutdown of glutamine biosynthesis from NH_3_ and glutamate abolishes NH_3_-dependent regulation of N_2_-fixation in genetically diverse bacteria. Targeted GS shutdown therefore affects NH_3_ excretion on two fronts, allowing sustained nitrogenase expression and activity in the presence of fixed N_2_ and preventing assimilation of NH_3_, favouring excretion into the environment.

Without establishing control of GS shutdown, engineered NH_3_ excreting diazotrophs are typically auxotrophic for glutamine, which would render them non-competitive in the environment (13, 16). Here, we placed expression of the *uAT-Ac2* allele under control of the NifA-inducible nitrogenase promoter P*nifH* which, when tuned correctly, triggered GS shutdown and NH_3_ excretion specifically under N_2_-fixing conditions. In the field, this could allow bacteria to retain competitiveness prior to forming O_2_-deplete biofilms on the surface of roots (63), however lack of host-specific control could permit provision of NH_3_ to non-target plant species. Thus, we further modified the engineered strain *Ac*PU-R22 by deleting *nifA* and bringing the mutant *nifA*_*L94Q/D95Q*_ and *rpoN* alleles under rhizopine-inducible control, permitting *in vitro* rhizopine-dependent activation of nitrogenase activity, GS shutdown and NH_3_ excretion. The resulting strain should also only activate these processes when colonising the roots or rhizosphere of transgenic *RhiP* barley (47), though we acknowledge that experimental demonstration of this will first require optimisation of the current rhizopine signalling circuitry that permits perception of rhizopine by 10-25% of cells colonising *RhiP* barley roots and activation of nitrogenase activity equating to approximately 15% of that observed for wild-type *Ac*LP cells colonising wild-type barley (46). We have made progress on improving rhizopine signalling here as our new low-copy rhizopine receiver plasmid pSIR03 permitted rhizopine-responsive GFP activation in over 90% of cells in the population, compared to our previously reported plasmid pSIN02 where only 36% of cells perceived rhizopine *in vitro* (46). Moving forward, increasing the proportion of *Ac* cells sensing rhizopine on the root surface and rhizosphere will be crucial to establish effective control of traits.

With bacteria engaging in partner-specific activation of NH_3_ excretion, there still exists the possibility that viability and competitiveness might suffer due to increased energy demand (14). Rhizobia overcome this problem by engaging in stringent signalling with the legume that permits partner-specific infection of nodules (64). Inside the nodule, the bacteria are provided with low-oxygen conditions conducive to nitrogenase stability, they can escape the fierce competition of the rhizosphere, and are fed carbon in the form of dicarboxylates (50, 51). Engineering a nodule-like niche with stringent entry requirements into cereals will likely be important to maximise the effectiveness of inoculation with engineered NH_3_ excreting inoculants. The strains developed here could be adapted for entry of such an environment and therefore, this work represents significant advancement towards the development of both associative and more intimate “synthetic N_2_-fixing symbiosis” with cereals.

## Materials and methods

### Bacterial strains and plasmids

Bacteria used in this study (S1 File) were cultured in TY (65) or UMS (66, 67) media supplemented with 300 μM nicotinic acid and 20 mM succinate as previously described (46). Plasmids (S2 Table) were constructed using HiFi assembly (New England Biolabs) or BEVA modular golden-gate assembly (68, 69) as outlined in the S1 File and were mobilised into *Azorhizobium* by diparental mating with *E. coli* ST18 (70). For mini-Tn*7* integration into the chromosome, tri-parental matings were used to additionally mobilise the transposase helper plasmid pTNS3, which carries an R6K origin of replication that is not maintained in *Azorhizobium* (71).

Gene deletion and replacement mutant strains were constructed by mobilising the relevant suicide plasmid, derived from pK19mobSacB (S2 Table & S1 File), into the target strain and selecting for single-crossover integration into the chromosomal region of interest by plating cells on selective UMS or TY agar media supplemented with 100 μg mL^−1^ kanamycin. Single-crossover mutants were subsequently grown in non-selective media until stationary phase and plated in serial dilutions onto UMS or TY agar supplemented with 10% (v/v) sucrose to select for double crossover deletion or replacement of the target gene. For the *ΔglnK*::Ω*Sp* replacement plasmid pOPS1564 only, 100 μg mL^−1^ spectinomycin and 1 mM IPTG was added to the media unless otherwise stated. Single colonies were patched onto the same media used for double-crossover selection plus and minus 100 μg mL^−1^ kanamycin and kanamycin sensitive colonies were screened by PCR and sanger sequencing for deletion or replacement of the target gene.

All *Ac*LP *ΔglnB ΔglnK*::Ω*Sp* mutant strains were constructed by first deleting *glnB* from *Ac*LP using plasmid pOPS1691, then subsequently integrating the *ΔglnK*::Ω*Sp* replacement plasmid pOPS1564 into the target chromosomal region by single-crossover. Because replacement of Δ*glnK*::Ω*Sp* was not possible on three separate occasions, mini-Tn*7* delivery plasmids carrying an IPTG-derepressible copy of *glnB* (S1 File) were integrated into the engineered *attB* site prior to selecting for selecting for double-crossover replacement of *glnK* with the Ω*Sp* interposon as described above.

### Growth curves

Growth curves were performed in triplicate by streaking single colonies onto 10 mL TY agar slopes and incubating for 3-days prior to three washes in PBS and inoculation at OD600λnm 0.01 into 500 μL UMS media in 24-well plates. The OD600λnm was monitored at 20 min intervals in an Omega FLUOstar plate reader set to shake cultures at 700 rpm at 37 °C until stationary phase. Growth statistics were calculated using the R package GrowthCurver (72).

### Glutamine synthetase transferase assays

Six-millilitre UMS cultures were initially grown in 30 mL glass universal vials sealed with silicone rubber septa as described for RT-qPCR experiments. After 3-h or 24-h incubation, 1 mL of culture was sampled for protein determination using a Millipore BCA protein assay kit. Five hundred microlitres of CTAB (1 mg mL^−1^) was added to the remaining cultures which were incubated at room temperature for a further 3 mins prior to harvesting by centrifugation at 4 °C. Cells were washed once with 5 mL 1% (w/v) KCL and finally resuspended in 500 μL of the same buffer and stored on ice. GS transferase assays were performed on 50 μL aliquots the permeabilised cells as previously described (15). The assays were performed in 500 μL total volumes with 30 min incubation in the presence or absence of 60 μM added Mg^2+^ to determine the total GS transferase activity and the activity of the “active” unadenylylated enzyme, respectively (48). The GS transferase buffer was adjusted to pH 7.0, as this was previously estimated as the iso-activity point for *Ac* (73). Following addition of the FeCl_3_ stop reagent, reaction tubes were centrifuged for 5 min at 13,000 g and 200 μL was transferred to clear, flat bottomed 96-well plates for spectrophotometric quantification of the product L-Glutamyl-γ-Hydroxamate (LGH) at 562λnm in a Promega GloMax multi-detection system.

### Acetylene reduction assays

Cultures for ARAs were prepared and analysed as previously described (46, 74) and 1 mL samples of the headspace atmosphere were analysed using a PerkinElmer Clarus 480 gas chromatograph equipped with a HayeSep® N (80–100 MESH) 584 column at 3-h, 5-h, 21-h, 23-h and 25-h incubation, unless otherwise stated.

### NH_3_ excretion assays

Three-millilitre UMS cultures were initially grown in 30 mL glass universal vials sealed with silicone rubber septa as described for RT-qPCR experiments. OD600λnm was recorded and NH_3_ was quantified in spent supernatants using the spectrophotometric indophenol assay as previously described (15). A calibration curve was performed for each experiment using freshly made dilutions of NH_3_Cl in UMS ranging from 5 μM – 1 mM. Absorbance of indophenol blue was quantified in a Genesys 150 UV visible spectrophotometer (Thermo Scientific) at 652λnm after 4-h incubation at room temperature.

### RT-qPCR

For RT-qPCR experiments, *n=*5 single colonies were streaked onto 10 mL UMS agar slopes supplemented with 20 mM succinate, 10 mM NH_4_Cl and 300 μM nicotinate and grown for 2-days at 37 °C. Cells were washed three times from the slopes with PBS, resuspended in UMS supplemented with the relevant carbon and N sources at OD600λnm 0.3 in 30 mL glass universal vials and transferred with the lid off into a sealed atmosphere cabinet adjusted to 3% O_2_ by flushing with N_2_ gas. After 30 min, cultures were sealed with silicone rubber septa and incubated at 37 °C with rigorous shaking for 3-h. Cells were next harvested by centrifugation at 4 °C, lysed using a FastPrep-24 5G instrument and cellular debris was removed by a second round of centrifugation. RNA was extracted from the resulting lysate using a Qiagen RNAeasy extraction kit and tested for quality and purity using an Agilent Experion Bioanalyzer with RNA Stdsens chips. gDNA was depleted from RNA by treatment with Invitrogen Turbo DNAse as per the manufacturer’s recommendations and 5 μg was used to generate cDNA using an Invitrogen SuperScript IV reverse transcriptase kit as per the manufacturer’s recommendations. The final cDNA template was diluted 1:20 with water and 1 μL was added to each 20 μL RT-qPCR reaction prepared in 96-well plates with Applied Biosystems PowerUp SYBR green master mix. Reactions were run using an Applied Biosystems ViiA 7 Real-Time PCR system. RT-qPCR primers were initially tested for amplification efficiency and target specificity by generating a standard curve of amplification with 5-fold dilutions of *Ac*LP gDNA. The housekeeping gene primer targeted *recA* and was validated previously (75), whereas the *glnA* primers designed here had the following sequence *glnA F* 5’-CCGCTGACCAACTCCTACA *glnA R* 5’-CCATGAACAGGGCCGAGAA.

### GFP reporter assays and flow-cytometry

GFP reporter assays and flow-cytometry experiments were performed on 24-h incubated cultures as previously described (46). Inducers were added directly to the growth media at the time of inoculation where relevant.

## Acknowledgements

The authors would like to thank Christian Rodgers for providing barley seeds for the experiments in this study.

## Supporting Information Captions

**S1 Fig. Characterisation of synthetic ribosome binding sites in *Ac*LP**. Each RBS was fused to GFP under expression by the strong synthetic promoter J23104 on plasmid pOGG024 and GFP was measured after 24-h incubation in UMS media (*n* = 3). Relative luminescence units are defined here as GFP fluorescence / OD600λnm. The RBS nucleotide sequences are provided in S1 File.

**S2 Fig. Expression and total activity of glutamine synthetase (*glnA*, GS) is elevated in *Ac*RGl. (a)** Total specific activity of both adenylated (inactive) and unadenylated (active) forms of GS was measured in whole cells grown for 24-h as determined by γ-glutamyl transferase assays (*n =* 5). **(b)** *glnA* expression was quantified relative to the housekeeping gene *recA* by RT-qPCR in cells growth for 3-h. All cultures for assays were grown in N_2_-fixing conditions (N-free UMS media with 3% O_2_ in the headspace). Error bars represent one SEM. Independent two-tailed students t-tests were used to compare means. ***P < 0.001.

**S3 Fig. Induction of the *Sinorhizobium meliloti* 1021 naringenin-inducible *PnodA* promoter in *Ac*LP (a)** Genetic schematic (not to scale) of the low-copy (RK2 replicon) naringenin-inducible *GFP* reporter plasmid pOPS1536. **(b)** GFP induction in *Ac*LP (*n =* 3) harbouring pOPSXXX in response to naringenin supplemented *in vitro*. Relative luminescence units are defined here as GFP fluorescence / OD600λnm.

**S4 Fig. Total activity of glutamine synthetase (GS) in** Δ***glnE* mutants expressing unidirectional adenylyl transferases from the non-induced P*nodA* promoter (a)** Total specific activity of both adenylated (inactive) and unadenylated (active) forms of GS was measured in whole cells grown in N_2_-fixing conditions (N-free UMS media with 3% O_2_ in the headspace) for 3-h as determined by γ-glutamyl transferase assays (*n =* 5 for wild-type *Ac*LP or *n* = 3 for other strains). Error bars represent one SEM. Independent two-tailed students t-tests were used to compare means against the wild-type (WT) *Ac*LP as a reference. Not significant (ns) indicates P > 0.05, *P < 0.05.

**S5 Fig. NH**_**3**_ **excretion is sub-optimal in** Δ***glnE* mutants expressing unidirectional adenylyl transferases from the P*nodA* promoter induced with naringenin**. Spectrophotometric determination of NH_3_ in media of cultures induced with 5 μM naringenin grown for 24-h in N_2_-fixing conditions (N-free UMS media with 3% O_2_ in the headspace). Error bars represent one SEM. *n =* 3 for wild-type *Ac*LP Δ*glnE* or *n* = 6 for other strains.

**S6 Fig. Growth statistics for control strains and** Δ***glnE* mutants expressing unidirectional adenylyltransferases**. Mean generation times and the max OD600λnm (i.e. the carrying capacity, k) were calculated from standard curves of cultures grown in UMS media at 21% O_2_. Representative growth curves are provided in Fig 3c and Fig 5b. Strains highlighted in white are wild-type (WT) *Ac*LP and *Ac*LPΔ*glnE* controls, strains highlighted in pink are *Ac*LPΔ*glnE* carrying P*nodA* [RBS] uAT-DT16 modules on parent plasmid pOGG093 and strains highlighted in blue are *Ac*LPΔ*glnE* carrying mini-Tn*7* integrated P*nifH* [RBS] uAT-*Ac2*-DT16 modules.

**S7 Fig. Rhizopine control of N**_**2**_ **fixation alone does not permit NH**_**3**_ **excretion**. Spectrophotometric determination of NH_3_ in media of *n =* 3 cultures grown for 24-h in N_2_-fixing conditions. Error bars represent one SEM. Strain *Azospirillum brasilense* HM053 was used here as a positive control.

**S8 Fig. NifA**_**L94Q/D95Q**_ **activity is tolerant to ambient environmental oxygen. (a)** Genetic schematic (not to scale) of the rhizopine *nifA*_*L94Q/D95Q*_-*rpoN* controller plasmid with P*nifH*::*GFP* reporter fusion pSIN03. **(b)** P*nifH* promoter activity was measured in *n =* 4 cultures grown for 24-h under the conditions indicated. Relative fluorescence units (RFU) are defined here as GFP fluorescence / OD600λnm. Error bars represent one SEM. Independent two-tailed students t-tests with Bonferroni-holm adjustment were used to compare means. P > 0.05. **P < 0.01, ***P < 0.001.

**S1 Table. Flowcytometry statistics for rhizopine-inducible GFP expression in *Ac*LP carrying pOPS1052**

**S2 Table. Plasmids used in this study**

## References

1. Awika JM. Major Cereal Grains Production and Use around the World. Advances in Cereal Science: Implications to Food Processing and Health Promotion. ACS Symposium Series. 1089: American Chemical Society; 2011. p. 1–13.

2. Udvardi M, Brodie EL, Riley W, Kaeppler S, Lynch J. Impacts of agricultural nitrogen on the environment and strategies to reduce these impacts. Procedia Environ Sci. 2015;29:303.

3. Bonilla Cedrez C, Chamberlin J, Guo Z, Hijmans RJ. Spatial variation in fertilizer prices in Sub-Saharan Africa. PLOS ONE. 2020;15(1):e0227764.

4. Holden ST. Fertilizer and sustainable intensification in Sub-Saharan Africa. Glob Food Sec. 2018;18:20–6.

5. Liu H, Carvalhais LC, Crawford M, Singh E, Dennis PG, Pieterse CMJ, et al. Inner plant values: diversity, colonization and benefits from endophytic bacteria. Front Microbiol. 2017;8:2552.

6. Souza Rd, Ambrosini A, Passaglia LMP. Plant growth-promoting bacteria as inoculants in agricultural soils. Genet Mol Biol. 2015;38(4):401–19.

7. Knights HE, Jorrin B, Haskett TL, Poole PS. Deciphering bacterial mechanisms of root colonization. Environmental Microbiology Reports. 2021;13(4):428–44.

8. Herridge DF, Peoples MB, Boddey RM. Global inputs of biological nitrogen fixation in agricultural systems. Plant Soil. 2008;311(1):1–18.

9. Haskett TL, Tkacz A, Poole PS. Engineering rhizobacteria for sustainable agriculture. ISME J. 2020;15:949–64.

10. Pedrosa FO, Oliveira ALM, Guimarães VF, Etto RM, Souza EM, Furmam FG, et al. The ammonium excreting Azospirillum brasilense strain HM053: a new alternative inoculant for maize. Plant Soil. 2020;451(1):45–56.

11. Dobbelaere S, Croonenborghs A, Thys A, Ptacek D, Vanderleyden J, Dutto P, et al. Responses of agronomically important crops to inoculation with Azospirillum. Funct Plant Biol. 2001;28(9):871–9.

12. Díaz-Zorita M, Fernández-Canigia MV. Field performance of a liquid formulation of Azospirillum brasilense on dryland wheat productivity. Eur J Soil Biol. 2009;45(1):3–11.

13. Bueno Batista M, Dixon R. Manipulating nitrogen regulation in diazotrophic bacteria for agronomic benefit. Biochem Soc Trans. 2019;47(2):603–14.

14. Inomura K, Bragg J, Follows MJ. A quantitative analysis of the direct and indirect costs of nitrogen fixation: a model based on Azotobacter vinelandii. ISME J. 2017;11(1):166–75.

15. Bueno Batista M, Brett P, Appia-Ayme C, Wang Y-P, Dixon R. Disrupting hierarchical control of nitrogen fixation enables carbon-dependent regulation of ammonia excretion in soil diazotrophs. PLoS Genet. 2021;17(6):e1009617.

16. Colnaghi R, Green A, He L, Rudnick P, Kennedy C. Strategies for increased ammonium production in free-living or plant associated nitrogen fixing bacteria. Plant Soil. 1997;194(1):145–54.

17. Bali A, Blanco G, Hill S, Kennedy C. Excretion of ammonium by a nifL mutant of Azotobacter vinelandii fixing nitrogen. Applied Environmental Microbiology. 1992;58(5):1711–8.

18. Brewin B, Woodley P, Drummond M. The basis of ammonium release in nifL mutants of Azotobacter vinelandii. J Bacteriol. 1999;181(23):7356–62.

19. Barney BM, Eberhart LJ, Ohlert JM, Knutson CM, Plunkett MH. Gene deletions resulting in increased nitrogen release by Azotobacter vinelandii: application of a novel nitrogen biosensor. Appl Environ Microbiol. 2015;81(13):4316–28.

20. Mus F, Khokhani D, MacIntyre AM, Rugoli E, Dixon R, Ané J-M, et al. Genetic determinants of ammonium excretion in nifL mutants of Azotobacter vinelandii. Appl Environ Microbiol. 2022;0(ja):AEM.01876-21.

21. Martinez-Argudo I, Little R, Dixon R. Role of the amino-terminal GAF domain of the NifA activator in controlling the response to the antiactivator protein NifL. Mol Microbiol. 2004;52(6):1731–44.

22. Reyes-Ramirez F, Little R, Dixon R. Mutant forms of the Azotobacter vinelandii transcriptional activator NifA resistant to inhibition by the NifL regulatory protein. J Bacteriol. 2002;184(24):6777.

23. Ghenov F, Gerhardt ECM, Huergo LF, Pedrosa FO, Wassem R, Souza EM. Characterization of glutamine synthetase from the ammonium-excreting strain HM053 of Azospirillum brasilense. Braz J Biol. 2021;82:e235927.

24. Machado HB, Funayama S, Rigo LU, Pedrosa FO. Excretion of ammonium by Azospirillum brasilense mutants resistant to ethylenediamine. Can J Microbiol. 1991;37(7):549–53.

25. Michel-Reydellet N, Desnoues N, Elmerich C, Kaminski PA. Characterization of Azorhizobium caulinodans glnB and glnA genes: involvement of the P_II_ protein in symbiotic nitrogen fixation. J Bacteriol. 1997;179(11):3580–7.

26. Toukdarian A, Saunders G, Selman-Sosa G, Santero E, Woodley P, Kennedy C. Molecular analysis of the Azotobacter vinelandii glnA gene encoding glutamine synthetase. J Bacteriol. 1990;172(11):6529–39.

27. Ambrosio R, Ortiz-Marquez JCF, Curatti L. Metabolic engineering of a diazotrophic bacterium improves ammonium release and biofertilization of plants and microalgae. Metab Eng. 2017;40:59–68.

28. Michel-Reydellet N, Kaminski PA. Azorhizobium caulinodans P_II_ and GlnK proteins control nitrogen fixation and ammonia assimilation. J Bacteriol. 1999;181(8):2655–8.

29. Schnabel T, Sattely E. Engineering posttranslational regulation of glutamine synthetase for controllable ammonia production in the plant symbiont Azospirillum brasilense. Applied Environmental Microbiology. 2021;87(14):e0058221.

30. Stadtman ER. Regulation of glutamine synthetase activity. EcoSal Plus. 2004;1(1).

31. Jaggi R, van Heeswijk WC, Westerhoff HV, Ollis DL, Vasudevan SG. The two opposing activities of adenylyl transferase reside in distinct homologous domains, with intramolecular signal transduction. EMBO J. 1997;16(18):5562–71.

32. Huergo LF, Chandra G, Merrick M. PII signal transduction proteins: nitrogen regulation and beyond. FEMS Microbiol Rev. 2013;37(2):251–83.

33. Kennedy C, Doetsch N, Meletzus D, Patriarca E, Amar M, Iaccarino M. Ammonium sensing in nitrogen fixing bacteria: Functions of the glnB and glnD gene products. Plant Soil. 1994;161(1):43–57.

34. Liang YY, de Zamaroczy M, Arséne F, Paquelin A, Elmerich C. Regulation of nitrogen fixation in Azospirillum brasilense Sp7: Involvement of nifA, glnA and glnB gene products. FEMS Microbiol Lett. 1992;100(1-3):113–9.

35. de Zamaroczy M. Structural homologues P(II) and P(Z) of Azospirillum brasilense provide intracellular signalling for selective regulation of various nitrogen-dependent functions. Mol Microbiol. 1998;29(2):449–63.

36. Meletzus D, Rudnick P, Doetsch N, Green A, Kennedy C. Characterization of the glnK-amtB operon of Azotobacter vinelandii. J Bacteriol. 1998;180(12):3260–4.

37. Hanson TE, Forchhammer K, de Marsac NT, Meeks JC. Characterization of the glnB gene product of Nostoc punctiforme strain ATCC 29133: glnB or the PII protein may be essential. Microbiology. 1998;144(6):1537–47.

38. Jonsson A, Nordlund S, Teixeira PF. Reduced activity of glutamine synthetase in Rhodospirillum rubrum mutants lacking the adenylyltransferase GlnE. Res Microbiol. 2009;160(8):581–4.

39. Mus F, Tseng A, Dixon R, Peters JW, Pettinari MJ. Diazotrophic growth allows Azotobacter vinelandii to overcome the deleterious effects of a glnE deletion. Appl Environ Microbiol. 2017;83(13):e00808–17.

40. Foor F, Janssen KA, Magasanik B. Regulation of synthesis of glutamine synthetase by adenylylated glutamine synthetase. Proc Natl Acad Sci. 1975;72(12):4844–8.

41. Parish T, Stoker NG. glnE is an essential gene in Mycobacterium tuberculosis. J Bacteriol. 2000;182(20):5715–20.

42. Carroll P, Pashley CA, Parish T. Functional analysis of GlnE, an essential adenylyl transferase in Mycobacterium tuberculosis. J Bacteriol. 2008;190(14):4894–902.

43. Schnabel T, Sattely E. Improved stability of engineered ammonia production in the plant-symbiont Azospirillum brasilense. ACS Synth Biol. 2021;10(11):2982–96.

44. Ryu MH, Zhang J, Toth T, Khokhani D, Geddes BA, Mus F, et al. Control of nitrogen fixation in bacteria that associate with cereals. Nat Microbiol. 2020;5(2):314–30.

45. Geddes BA, Ryu MH, Mus F, Garcia Costas A, Peters JW, Voigt CA, et al. Use of plant colonizing bacteria as chassis for transfer of N_2_-fixation to cereals. Curr Opin Biotechnol. 2015;32:216–22.

46. Haskett TL, Paramasivan P, Mendes MD, Green P, Geddes B, Knights HE, et al. Engineered plant control of associative nitrogen fixation. Proc Natl Acad Sci. 2022;in press.

47. Geddes BA, Paramasivan P, Joffrin A, Thompson AL, Christensen K, Jorrin B, et al. Engineering transkingdom signalling in plants to control gene expression in rhizosphere bacteria. Nat Commun. 2019;10(1):3430.

48. Bender RA, Janssen KA, Resnick AD, Blumenberg M, Foor F, Magasanik B. Biochemical parameters of glutamine synthetase from Klebsiella aerogenes. J Bacteriol. 1977;129(2):1001–9.

49. Bolleter WT, Bushman CJ, Tidwell PW. Spectrophotometric Determination of Ammonia as Indophenol. Analytical Chemistry. 1961;33(4):592–4.

50. Schulte CCM, Borah K, Wheatley RM, Terpolilli JJ, Saalbach G, Crang N, et al. Metabolic control of nitrogen fixation in rhizobium-legume symbioses. Science Advances. 2021;7(31).

51. Rutten PJ, Steel H, Hood GA, Ramachandran VK, McMurtry L, Geddes B, et al. Multiple sensors provide spatiotemporal oxygen regulation of gene expression in a Rhizobium-legume symbiosis. PLoS Genet. 2021;17(2):e1009099.

52. Fischer H-M, Hennecke H. Direct response of Bradyrhizobium japonicum nifA-mediated nif gene regulation to cellular oxygen status. Mol Gen Genet. 1987;209(3):621–6.

53. Michel-Reydellet N, Desnoues N, de Zamaroczy M, Elmerich C, Kaminski PA. Characterisation of the glnK-amtB operon and the involvement of AmtB in methylammonium uptake in Azorhizobium caulinodans. Mol Gen Genet. 1998;258(6):671–7.

54. Drepper T, Gross S, Yakunin AF, Hallenbeck PC, Masepohl B, Klipp W. Role of GlnB and GlnK in ammonium control of both nitrogenase systems in the phototrophic bacterium Rhodobacter capsulatus. Microbiology. 2003;149(Pt 8):2203–12.

55. Pekgöz G, Gündüz U, Eroğlu I, Yücel M, Kovács K, Rákhely G. Effect of inactivation of genes involved in ammonium regulation on the biohydrogen production of Rhodobacter capsulatus. International Journal of Hydrogen Energy. 2011;36(21):13536–46.

56. Neilson AH, Nordlund S. Regulation of nitrogenase synthesis in intact cells of Rhodospirillum rubrum: inactivation of nitrogen fixation by ammonia, L-glutamine and L-asparagine. J Gen Microbiol. 1975;91(1):53–62.

57. Turpin DH, Edie SA, Canvin DT. In vivo nitrogenase regulation by ammonium and methylamine and the effect of MSX on ammonium transport in Anabaena flos-aquae. Plant Physiol. 1984;74(3):701–4.

58. Reich S, Almon H, Böger P. Short-term effect of ammonia on nitrogenase activity of Anabaena variabilis (ATCC29413). FEMS Microbiol Lett. 1986;34(1):53–6.

59. Jones BL, Monty KJ. Glutamine as a feedback inhibitor of the Rhodopseudomonas sphaeroides nitrogenase system. J Bacteriol. 1979;139(3):1007–13.

60. Kuczius T, Kleiner D. Ammonia-excreting mutants of Klebsiella pneumoniae with a pleiotropic defect in nitrogen metabolism. Arch Microbiol. 1996;166(6):388–93.

61. Masepohl B, Drepper T, Paschen A, Gross S, Pawlowski A, Raabe K, et al. Regulation of nitrogen fixation in the phototrophic purple bacterium Rhodobacter capsulatus. J Mol Microbiol Biotechnol. 2002;4(3):243–8.

62. Masepohl B, Klipp W. Organization and regulation of genes encoding the molybdenum nitrogenase and the alternative nitrogenase in Rhodobacter capsulatus. Arch Microbiol. 1996;165(2):80–90.

63. Wang D, Xu A, Elmerich C, Ma LZ. Biofilm formation enables free-living nitrogen-fixing rhizobacteria to fix nitrogen under aerobic conditions. ISME J. 2017;11(7):1602–13.

64. Bozsoki Z, Gysel K, Hansen SB, Lironi D, Krönauer C, Feng F, et al. Ligand-recognizing motifs in plant LysM receptors are major determinants of specificity. Science. 2020;369(6504):663–70.

65. Beringer JE. R factor transfer in Rhizobium leguminosarum. Microbiology. 1974;84(1):188–98.

66. Poole PS, Schofield NA, Reid CJ, Drew EM, Walshaw DL. Identification of chromosomal genes located downstream of dctD that affect the requirement for calcium and the lipopolysaccharide layer of Rhizobium leguminosarum. Microbiology. 1994;140 (Pt 10):2797–809.

67. Brown CM, Dilworth MJ. Ammonia assimilation by Rhizobium cultures and bacteroids. Microbiology. 1975;86(1):39–48.

68. Geddes BA, Mendoza-Suárez MA, Poole PS. A bacterial expression vector archive (BEVA) for flexible modular assembly of golden gate-compatible vectors. Front Microbiol. 2019;9:3345.

69. Weber E, Engler C, Gruetzner R, Werner S, Marillonnet S. A modular cloning system for standardized assembly of multigene constructs. PLoS One. 2011;6(2):e16765.

70. Thoma S, Schobert M. An improved Escherichia coli donor strain for diparental mating. FEMS Microbiol Lett. 2009;294(2):127–32.

71. Choi KH, Schweizer HP. mini-Tn7 insertion in bacteria with single attTn7 sites: example Pseudomonas aeruginosa. Nat Protoc. 2006;1(1):153–61.

72. Sprouffske K, Wagner A. Growthcurver: an R package for obtaining interpretable metrics from microbial growth curves. BMC Bioinformatics. 2016;17(1):172.

73. Donald RG, Ludwig RA. Rhizobium sp. strain ORS571 ammonium assimilation and nitrogen fixation. J Bacteriol. 1984;158(3):1144–51.

74. Haskett TL, Knights HE, Jorrin B, Mendes MD, Poole PS. A simple in situ assay to assess plant-associative bacterial nitrogenase activity. Front Microbiol. 2021;12(1598).

75. Ling J, Wang H, Wu P, Li T, Tang Y, Naseer N, et al. Plant nodulation inducers enhance horizontal gene transfer of Azorhizobium caulinodans symbiosis island. Proc Natl Acad Sci. 2016;113(48):13875–80.

