## Supplementary material for "Control of N_2_ fixation and NH_3_ excretion in *Azorhizobium caulinodans* ORS571": S1 Table

| **Sample** | ***Count mCherry+** | **†Count GFP+** | **Count GFP**  **(% of mCherry+)** |
| --- | --- | --- | --- |
| Not induced | 13910 | 1 | 7.19E-03 |
| Not induced | 13804 | 1 | 7.24E-03 |
| Not induced | 13582 | 0 | 0 |
| Not induced | 13960 | 1 | 7.16E-03 |
| + 10 uM SI | 12651 | 11675 | 92.3 |
| + 10 uM SI | 11247 | 10443 | 92.9 |
| + 10 uM SI | 12738 | 11844 | 93 |
| + 10 uM SI | 12859 | 12104 | 94.1 |

***** mCherry+ is defined as bacteria where mCherry fluorescence was above 3000 a.u.

**†** GFP+ is defined here as bacteria where GFP fluorescence was above the mean 99^th^ percentile of the uninduced state.
