## Supplementary material for "Control of N_2_ fixation and NH_3_ excretion in *Azorhizobium caulinodans* ORS571": S2 Table

| **Plasmid** | **Replicon** | **Antibiotic resistance** | **Description** | **Ref** |
| --- | --- | --- | --- | --- |
| pSRK-Km | pBBR1 | Kan | Broad-host-range cloning vector with IPTG-derepressible promoter | (1) |
| pOPS1691 | pBR322 | Kan | *glnB* markerless deletion vector | This study |
| pOPS1855 | pBR322 | Kan | *glnE* markerless deletion vector | This study |
| pOPS1564 | pBR322 | Kan,Spec | *glnK* markerless deletion vector | This study |
| pOPS1563 | pBR322 | Kan,Spec | *glnK* omega-spectinomycin cassette replacement vector | This study |
| pOGG276 | R6K | Gent,Carb | Low-copy mini-Tn*7* golden-gate destination vector | B. Jorrin |
| pOPS1531 | R6K | Gent,Carb | mini-Tn*7* delivery plasmid with J23104-[Rstd]-*mCherry-*DT16 cassette | (2) |
| pOPS1638 | R6K | Gent | mini-Tn7 delivery plasmid with NifA-inducible *lacI* with RBS [R22] and LacI-repressible *glnB* with RBS [R1] | This study |
| pOPS1640 | R6K | Gent | mini-Tn7 delivery plasmid with NifA-inducible *lacI* with RBS [R22] and LacI-repressible *glnB* with RBS [R28] | This study |
| pOPS1639 | R6K | Gent | mini-Tn7 delivery plasmid with NifA-inducible *lacI* with RBS [R22] and LacI-repressible *glnB* with RBS [R31] | This study |
| pOPS1784 | R6K | Gent,Carb | mini-Tn7 delivery plasmid with NifA-inducible *lacI* with RBS [R22] and LacI-repressible *glnB* with RBS [RStd] | This study |
| pOPS1878 | R6K | Gent,Carb | mini-Tn7 delivery plasmid with NifA-inducible uAT-Ac2 with RBS [native PnifH RBS] | This study |
| pOPS1874 | R6K | Gent,Carb | mini-Tn7 delivery plasmid with NifA-inducible uAT-Ac2 with RBS [R1] | This study |
| pOPS1875 | R6K | Gent,Carb | mini-Tn7 delivery plasmid with NifA-inducible uAT-Ac2 with RBS [R22] | This study |
| pOPS1876 | R6K | Gent,Carb | mini-Tn7 delivery plasmid with NifA-inducible uAT-Ac2 with RBS [R28] | This study |
| pOPS1877 | R6K | Gent,Carb | mini-Tn7 delivery plasmid with NifA-inducible uAT-Ac2 with RBS [R31] | This study |
| pOPS1873 | R6K | Gent,Carb | mini-Tn7 delivery plasmid with NifA-inducible uAT-Ac2 with RBS [Rstd] | This study |
| pOPS1536 | RK2 | Tet | Naringenin-inducible *gfp* | This study |
| pOPS1859 | RK2 | Tet | Naringenin-inducible uAT-Ac1 with RBS [Rstd] | This study |
| pOPS1860 | RK2 | Tet | Naringenin-inducible uAT-Ac2 with RBS [Rstd] | This study |
| pOPS1861 | RK2 | Tet | Naringenin-inducible uAT-Ac3 with RBS [Rstd] | This study |
| pOPS1857 | RK2 | Tet | Naringenin-inducible uAT-Ec10 with RBS [Rstd] | This study |
| pOPS1858 | RK2 | Tet | Naringenin-inducible uAT-Ec11 with RBS [Rstd] | This study |
| pOPS1565 | pBR322 | Kan | *nifA* markerless deletion vector | PNAS |
| pOPS1213 | pBBR1 | Gent | NifA-inducble *gfp* | (2) |
| pOGG037 | ColE1 | Spec | pL0M SC *GFP* Level 0 golden-gate SC module | (3) |
| pOGG120 | ColE1 | Spec | pL0M-P PJ23104 Level 0 golden-gate promoter module | (4) |
| pOGG324 | ColE1 | Spec | pL0M-SC *lacI* Level 0 golden-gate CDS module | This study |
| pOGG157 | ColE1 | Spec | pL0M-T DT16 Level 0 golden-gate terminator module | (4) |
| pOGG162 | ColE1 | Spec | pL0M-T F6S Level 0 golden-gate terminator module | (4) |
| pOGG144 | ColE1 | Spec | pL0M-U [R1] Level 0 golden-gate RBS module | (4) |
| pOGG145 | ColE1 | Spec | pL0M-U [R13] Level 0 golden-gate RBS module | (4) |
| pOGG146 | ColE1 | Spec | pL0M-U [R22] Level 0 golden-gate RBS module | (4) |
| pOGG271 | ColE1 | Spec | pL0M-U [R25] Level 0 golden-gate RBS module | This study |
| pOGG272 | ColE1 | Spec | pL0M-U [R28] Level 0 golden-gate RBS module | This study |
| pOGG147 | ColE1 | Spec | pL0M-U [R31] Level 0 golden-gate RBS module | (4) |
| pOGG143 | ColE1 | Spec | pL0M-U [RStd] Level 0 golden-gate RBS module | (4) |
| pOGG284 | RK2 | Carb | pL1M-F1 Level 1 golden-gate destination vector, very low copy number | This study |
| pOGG359 | RK2 | Carb | pL1M-F1 P*lac* [R1] *glnB* DT16 Level 1 golden-gate module | This study |
| pOGG361 | RK2 | Carb | pL1M-F1 P*lac* [R28] *glnB* DT16 Level 1 golden-gate module | This study |
| pOGG360 | RK2 | Carb | pL1M-F1 P*lac* [R31] *glnB* DT16 Level 1 golden-gate module | This study |
| pOGG308 | ColE1 | Carb | pL1M-F1 P*lac* [RStd] *glnB* DT16 Level 1 golden-gate module | This study |
| pOGG054 | ColE1/RK2 | Carb | pL1M-F2 Level 1 golden-gate destination vector, high copy number | (3) |
| pOGG322 | RK2 | Carb | pL1M-F2 Level 1 golden-gate destination vector, very low copy number | This study |
| pOPS1778 | pBR322 | Gent, Carb | pL1M-F2 Level 1 PnifH-[R22]-*lacI*-F6S | This study |
| pSIR03 | RK2 | Tc | Rhizopine-inducible *gfp* | This study |
| pSIN02 | pBBR1 | Kan | Rhizopine-inducible *nifA*(L94Q/D95Q)-*rpoN* cassette, medium copy number | (5) |
| pSIN03 | pBBR1 | Kan | Rhizopine-inducible *nifA*(L94Q/D95Q)-*rpoN* cassette with P*nifH::GFP* promoter fusion, medium copy number | (5) |
| pSIN04 | RK2 | Tet | Rhizopine-inducible *nifA*(L94Q/D95Q)-*rpoN* cassette, very low copy number | This study |
| pOGG093 | RK2 | Tet | Stable broad-host-range golden-gate destination vector, very low copy number | (3) |
| pK19mobsacB | pBR322 | Kan | Suicide vector for construction of markerless deletion mutants | (6) |
| pHP45ΩSp | pMB1 | Spec, Carb | Ω-Spectinomycin cloning vector | (7) |

1.	S. R. Khan, J. Gaines, R. M. Roop, 2nd, S. K. Farrand, Broad-host-range expression vectors with tightly regulated promoters and their use to examine the influence of TraR and TraM expression on Ti plasmid quorum sensing. *Appl. Environ. Microbiol.* **74**, 5053-5062 (2008).

2.	T. L. Haskett, H. E. Knights, B. Jorrin, M. D. Mendes, P. S. Poole, A simple in situ assay to assess plant-associative bacterial nitrogenase activity. *Front. Microbiol.* **12** (2021).

3.	B. A. Geddes, M. A. Mendoza-Suárez, P. S. Poole, A bacterial expression vector archive (BEVA) for flexible modular assembly of golden gate-compatible vectors. *Front. Microbiol.* **9**, 3345 (2019).

4.	K. Grant (2019) Engineering rhizobacteria as synthetic biology chassis. in *Department of Plant Sciences* (University of Oxford).

5.	T. L. Haskett *et al.*, Engineered plant control of associative nitrogen fixation. *Proc. Natl. Acad. Sci* ***in press*** (2022).

6.	A. Schäfer *et al.*, Small mobilizable multi-purpose cloning vectors derived from the *Escherichia coli* plasmids pK18 and pK19: selection of defined deletions in the chromosome of *Corynebacterium glutamicum*. *Gene* **145**, 69-73 (1994).

7.	P. Prentki, H. M. Krisch, *In vitro* insertional mutagenesis with a selectable DNA fragment. *Gene* **29**, 303-313 (1984).
