## Supplementary figures and images for "Control of N_2_ fixation and NH_3_ excretion in *Azorhizobium caulinodans* ORS571"

### S1 Fig

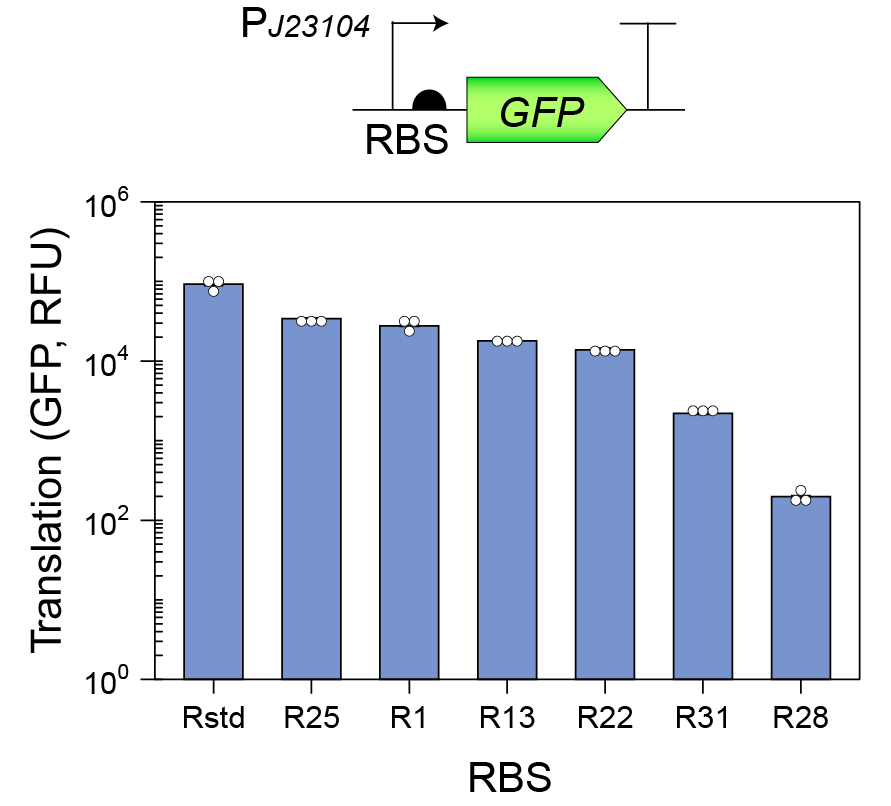

### S2 Fig

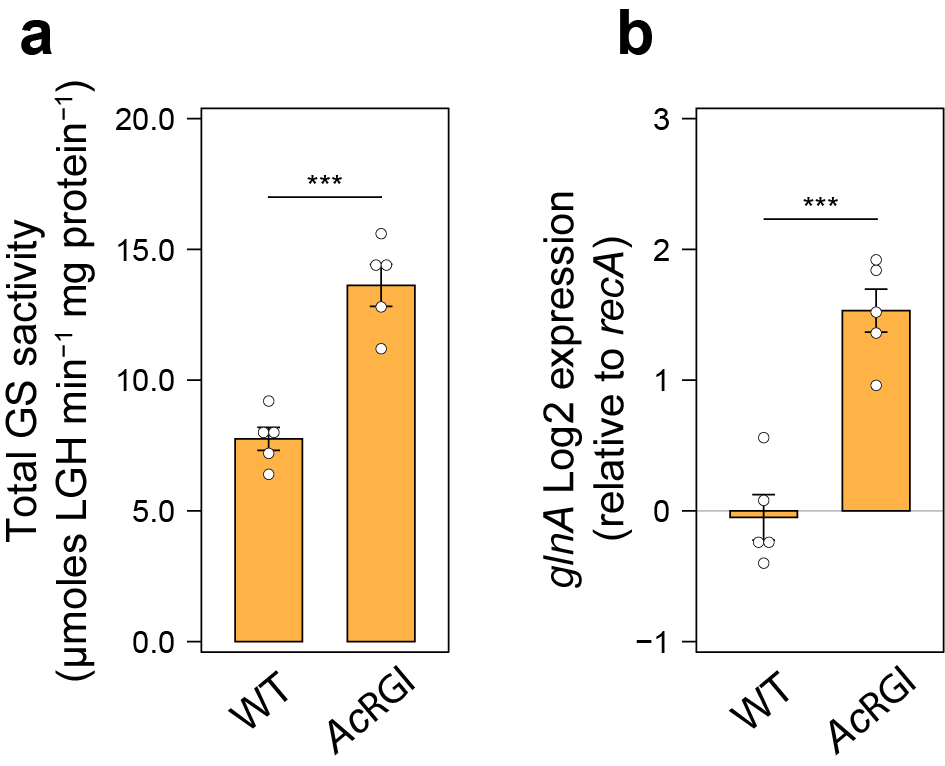

### S3 Fig

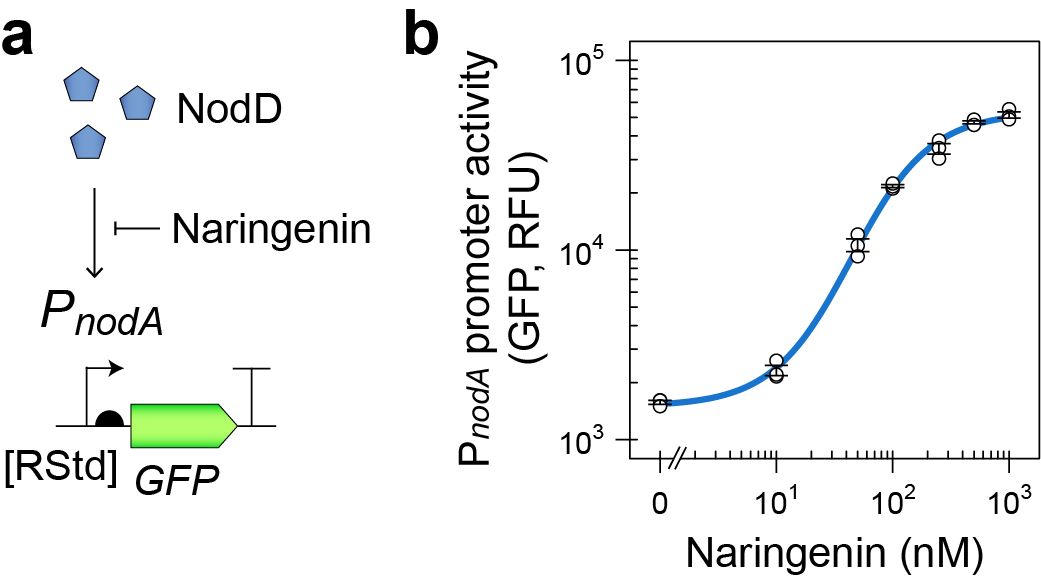

### S4 Fig

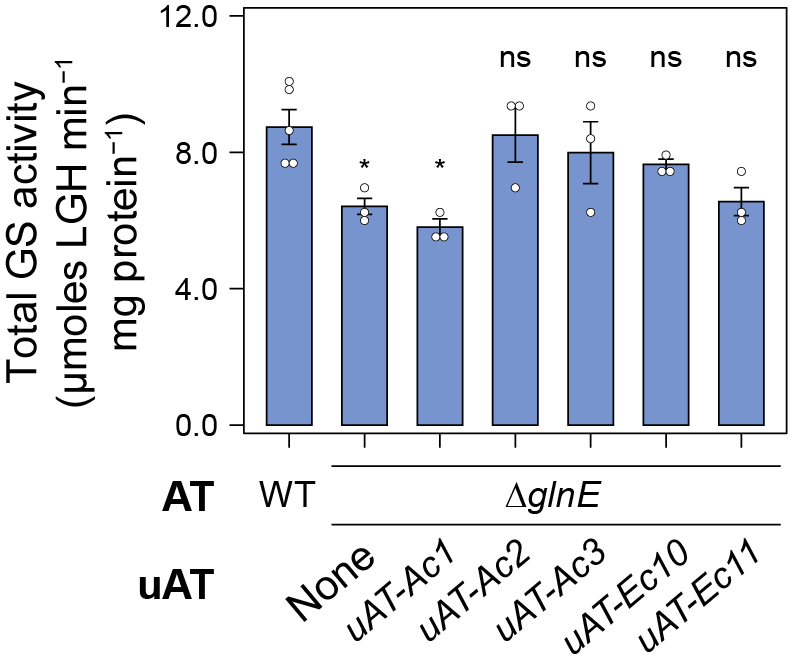

### S5 Fig

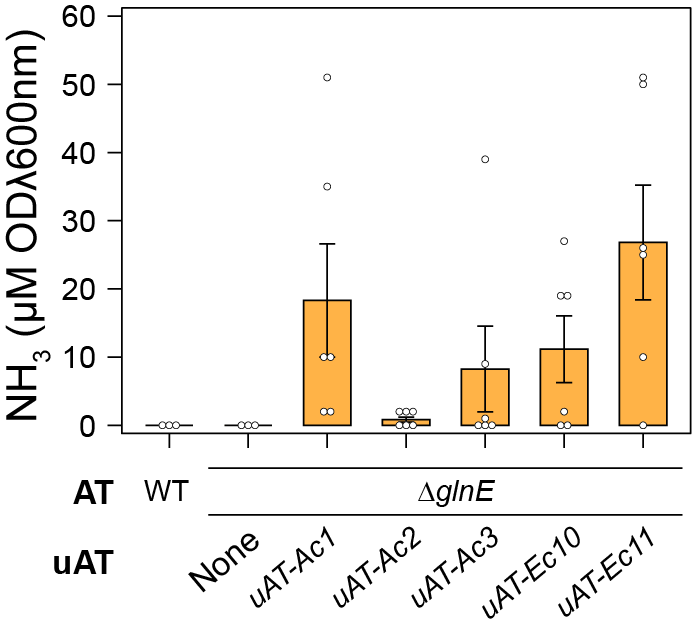

### S6 Fig

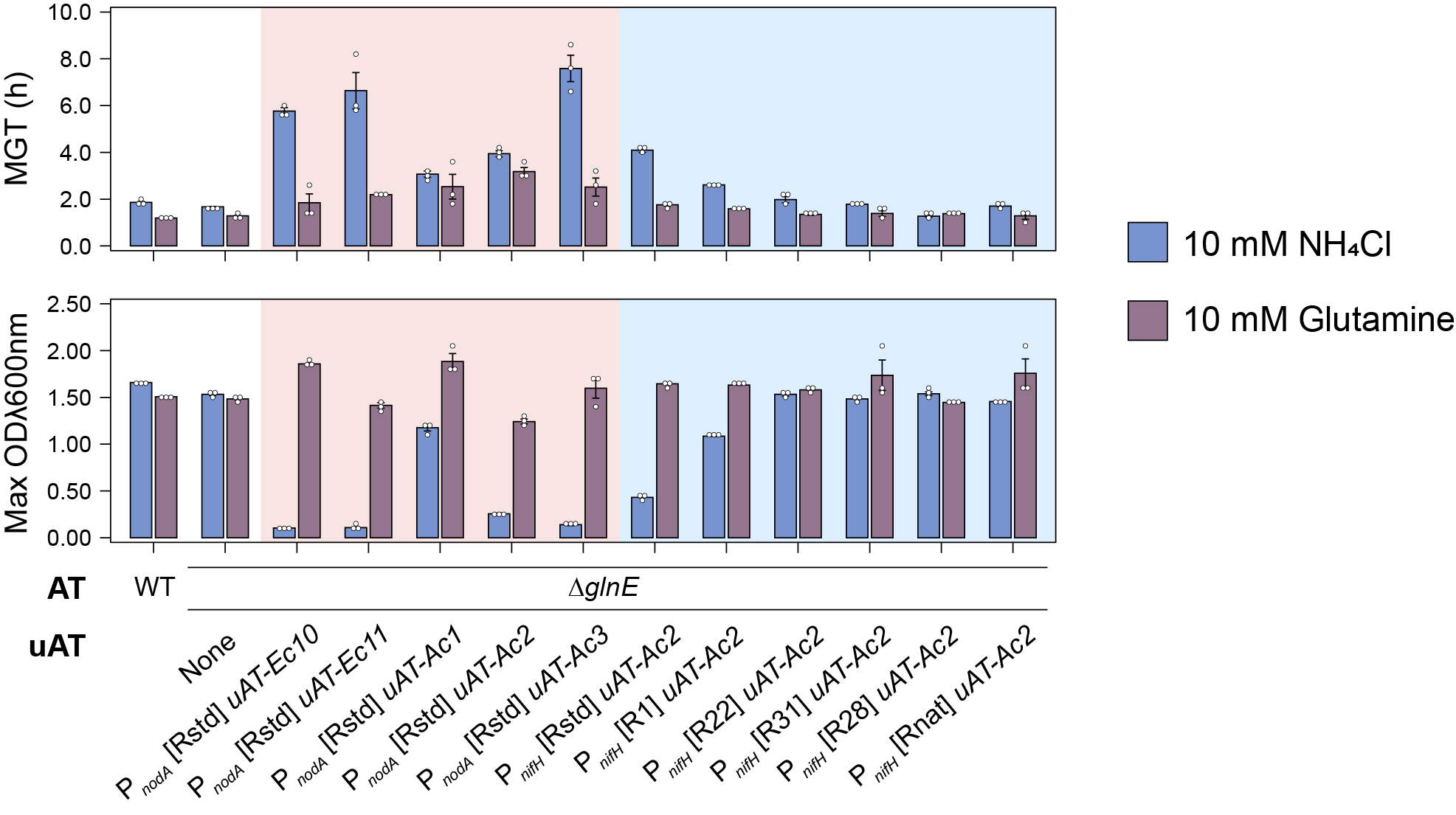

### S7 Fig

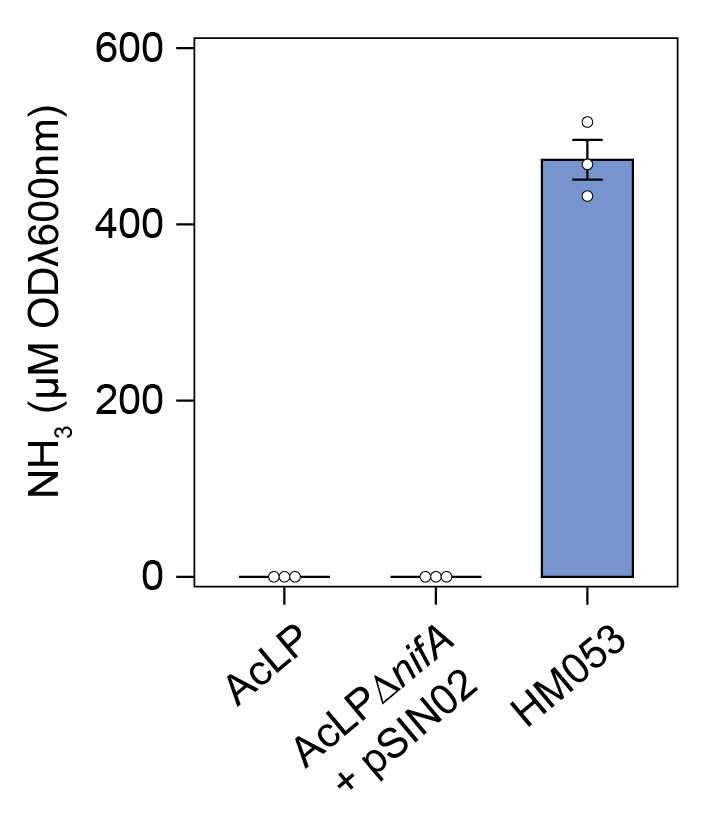

### S8 Fig

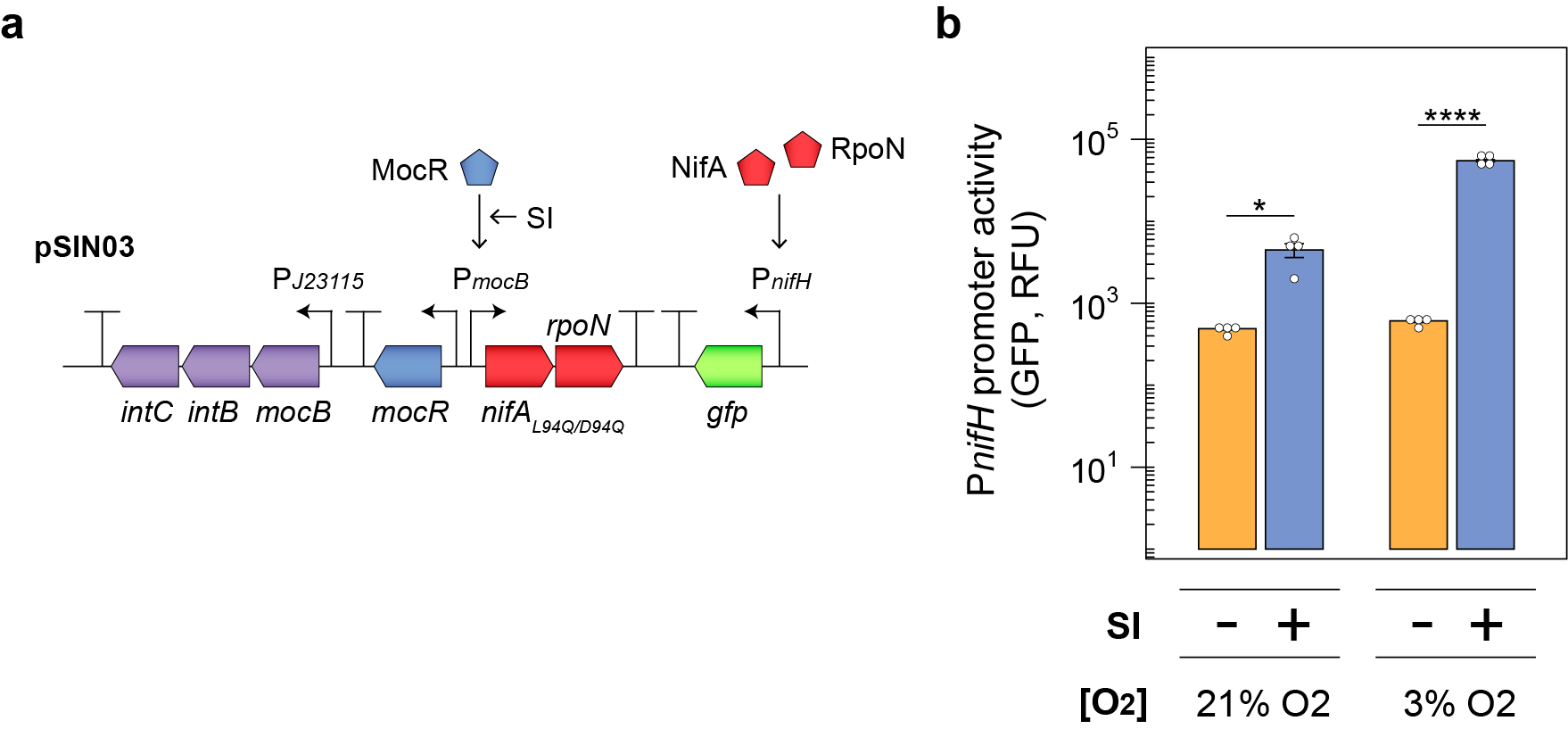
